# Locomotor Trends Among Early Mammals Illuminated by Predictors of Arboreality from the Mammalian Appendicular Skeleton

**DOI:** 10.1101/2025.07.09.663922

**Authors:** Jonathan A. Nations, Lucas N. Weaver, David M. Grossnickle

**Affiliations:** Florida Museum of Natural History, University of Florida, Gainesville, FL, 32610, USA; Museum of Paleontology & Department of Earth and Environmental Sciences, University of Michigan, Ann Arbor, MI, 48109, USA; Department of Earth and Environmental Sciences, University of Michigan, Ann Arbor, MI 48109, USA; Natural Sciences Department, Oregon Institute of Technology, Klamath Falls, OR, 97601, USA

**Keywords:** Functional morphology, Ecomorphology, Mesozoic, Therians, Multituberculates, Plesiadapiforms, Ordinal model, Bayesian

## Abstract

Arboreal locomotion is a common trait among extant mammals, and it likely evolved independently in numerous Mesozoic and Cenozoic mammalian lineages. We review evidence for the evolution of arboreality within lineages of early mammals and their stem relatives (Synapsida) while also emphasizing the uncertainty of locomotor inferences of fossil lineages. An improved understanding of the evolutionary origins and prevalence of arboreality in early mammals requires robust links between morphological traits and locomotor behaviors that can be applied to fossils. To this end, we use a large comparative dataset of linear measurements taken on extant mammalian postcranial specimens to evaluate the skeletal proxies that best differentiate arboreality from other locomotor modes. Consistent with previous research, we find that measures of digit length and elbow proportions are especially powerful predictors of climbing. Locomotor predictions based on single measurements or ratios are improved when phylogeny is incorporated into models. Nonetheless, combining multiple functionally relevant traits into our models yields robust predictions of locomotor mode even without incorporating phylogenetic information—this is especially encouraging for fossil groups with unstable phylogenies. We test these locomotor proxies in three early-mammalian case studies—Multituberculata, Plesiadapiformes, and early Theria. We find that multituberculates had a wide locomotor breadth whereas plesiadapiforms all had a high probability of climbing, consistent with previous studies. However, some Mesozoic therians in our sample are predicted to be ground-dwellers, conflicting with the hypothesis that the ancestral therian was scansorial or arboreal. The predictive tools presented here will improve locomotor inferences for small mammals, further illuminating the prevalence and importance of arboreality in early mammalian evolution.

## 1 Introduction

The locomotor diversity among extant mammals exceeds that of any other tetrapod group, spanning the range of fully aquatic (e.g., whales) to fully volant (i.e., bats) taxa and nearly everything in between (e.g., Nowak, 1999). Of these numerous locomotor modes, arboreality has been of special interest to researchers. This is due to several factors, such as arboreality (1) is an ecological trait that can often be inferred in fossil or poorly studied taxa via simple limb traits, (2) is thought to require morphological adaptations to cope with changes in substrate shape and gravitational force, (3) represents an ecological shift that has evolved repeatedly (and perhaps secondarily, see below) in nearly every “clade” of terrestrial mammals, (4) is hypothesized to be a key factor in the evolution of primates, and (5) may have been the ancestral locomotor mode of major mammalian groups, especially therian mammals (placentals and marsupials; e.g., Huxley, 1880; Dollo, 1899; Bensley, 1903; Matthew, 1904; Szalay, 1981; Szalay, 1984; Szalay and Sargis, 2001; Ji et al., 2002; Luo et al., 2003; O’Leary et al., 2013; Chen and Wilson, 2015; Bi et al., 2018). Because arboreality may be linked to the evolutionary origins of several major mammalian groups, it is often posited to have played a prominent role in mammalian evolution, and possibly to have co-evolved with major plant groups (e.g., angiosperm trees in the Late Cretaceous and/or Paleogene; Wing and Tiffney, 1987; Collinson and Hooker, 1991; Sussman 1991; Eriksson, 2016). Thus, investigating the evolution of mammalian arboreality is not only critical for understanding organismal-level information such as morphological adaptations and ecological niches, but it is also important for understanding broad mammalian macroevolutionary and macroecological patterns.

Our understanding of when temporally and where phylogenetically arboreality has arisen and its prevalence through mammalian evolution comes mostly from the fossil record, which, although limited to teeth and jaw fragments for many taxa, does yield postcranial specimens. For Mesozoic mammals, in particular, there has been an explosion of new fossil discoveries over the past 25 years that include postcranial material, revising our understanding of locomotor strategies and diversity in early mammals (Luo, 2007; Meng, 2014; Chen and Wilson, 2015; Grossnickle et al., 2019). These fossil specimens include lineages exhibiting evidence of arboreal/scansorial (Luo et al., 2011; Bi et al., 2014, Grossnickle et al., 2020), gliding (Han et al., 2017; Meng et al., 2017; Grossnickle et al., 2020), fossorial (Luo and Wible, 2005; Meng et al., 2015), and semi-aquatic (Ji et al., 2006) locomotion as early as the Jurassic Period. Critical to inferring the paleoecologies of fossil lineages is understanding how morphological traits are associated with specific behaviors, including how limb traits are linked to locomotor strategies. Thus, studying ecomorphological traits of extant mammals is important for deciphering the ecologies of early mammals.

Despite the expanding fossil record of postcranial skeletons, it remains challenging to infer locomotor strategies in early mammals. For instance, early mammalian fossils preserving postcrania are almost always fragmentary and, even in rare exceptions when most of the skeletal elements are preserved, early branching mammalian lineages often exhibit a suite of symplesiomorphic traits of non-mammalian cynodonts (hereafter ‘cynodonts’) or specialized apomorphies that confound attempts to infer locomotor strategies via comparisons with living taxa (e.g., Jenkins and Parrington, 1976; Panciroli et al., 2022). This uncertainty complicates our understanding of both when and where arboreality evolved and its importance in mammalian evolution.

The appendicular skeleton has long been used to infer the locomotor behaviors of both extant and extinct mammals (e.g., Huxley, 1880; Dollo, 1899; Bensley, 1903; Matthew, 1904; Maynard Smith and Savage, 1956; Haines, 1958; Jenkins, 1971; Hildebrand, 1985a; Hildebrand, 1985b; Szalay and Sargis, 2001; Elissamburu and Vizcaíno, 2004; Dunn, 2018). Though many studies have focused on primates, often such studies use non-primate taxa to illuminate possible evolutionary trajectories to arboreality (Sargis, 2002a; Sargis, 2002b; Gebo, 2004; Kirk et al., 2008). Additional research into mammal arboreality outside of primates has focused on inferring locomotion in extinct and poorly studied extant taxa (e.g., Haines, 1958; Chen and Wilson, 2015). Combined, these studies have identified a suite of postcranial measurements that may covary with locomotion in mammals, such as relative phalangeal dimensions and relative length of the ulnar olecranon process (Maynard Smith and Savage, 1956; Hildebrand, 1985a; Hildebrand, 1985b; Lemelin, 1999; Bloch and Boyer, 2002; Elissamburu and Vizcaíno, 2004; Polly, 2007; Kirk et al., 2008; Samuels and Van Valkenburgh, 2008; Samuels et al., 2013; Gould and Rose, 2014; Chen and Wilson, 2015; Kilbourne, 2017; Dunn, 2018; Nations et al., 2019; Weaver and Grossnickle, 2020; Burtner et al., 2024). Further, other studies have identified differences in shapes of articular surfaces among mammalian locomotor modes (e.g., Argot, 2001; Argot, 2002, Sargis, 2002a; Sargis, 2002b; Janis and Martin-Serra, 2020; Panciroli et al., 2022).

That early mammals were small-bodied (rarely exceeding 1 kg) is a truism that has complicated inferences about their locomotor diversity. A common concern in mammalian form-function studies is that small mammals may not exhibit distinct functional limb morphologies among locomotor modes (e.g., Hedrick et al., 2020). Jenkins (1974) argued that among small mammals, primarily ground-dwelling species require the same locomotor repertoire and associated postcranial adaptations as primarily tree-dwelling species to contend with uneven and disordered substrates (e.g., tree roots, woody debris). Jenkins and Parrington (1976) elaborated upon that hypothesis in their study of the postcranial anatomy and functional morphology of Triassic mammaliaforms (i.e., stem mammals), arguing that it was largely futile to ascribe a terrestrial vs. arboreal locomotor category to Mesozoic mammals, since they would all have had to climb to some degree. Jenkins’ hypothesis thus implies that the postcranial skeletons of small-bodied arborealists and terrestrialists should be nearly indistinguishable (Jenkins 1974; Jenkins and Parrington 1976), making inferences of locomotion in fossil small mammals especially challenging.

Nonetheless, mounting evidence indicates that Jenkins’ hypothesis may not hold true, with numerous studies on diverse mammalian subclades highlighting that small tree-dwelling and ground-dwelling mammals exhibit distinct morphologies that reflect their locomotor modes (Szalay, 1984; Argot, 2001; Szalay and Sargis, 2001, Argot, 2002; Sargis 2002a; Sargis, 2002b; Salton and Sargis, 2008; Salton and Sargis, 2009). Further, multivariate analyses of postcranial measurements have successfully differentiated small mammals on the basis of their locomotor modes (Samuels and Van Valkenburgh, 2008; Hopkins and Davis, 2009; Samuels et al., 2013; Chen and Wilson, 2015; Meng et al., 2017; Calede et al., 2019; Nations et al., 2019; Grossnickle et al., 2020; Janis and Martin-Serra, 2020; Weaver and Grossnickle, 2020). Finally, Weaver and Grossnickle (2020) directly tested predictions of Jenkins’ hypothesis and their analyses largely refuted the hypothesis. Although somewhat counterintuitive—small mammals do exhibit considerable locomotor plasticity (e.g., Nitikman and Mares, 1987; Granatosky, 2018) and have to contend with more complex substrates (e.g., August, 1983; Malcolm, 1995)—the postcranial morphologies of small mammals can be differentiated by locomotor modes to nearly the same extent as those of medium-sized mammals; that suggests small mammalian skeletons are adapted to avoid the most dangerous aspects of their locomotor environment (perhaps falling for tree-dwellers [e.g., Cartmill, 1985] and predation for ground-dwellers [e.g., Shattuck and Williams, 2010]), rather than to perfect a generalized locomotor repertoire (Weaver and Grossnickle, 2020). In sum, these mounting refutations of Jenkins’ hypothesis indicate that limb morphology is indeed a powerful predictor of locomotor mode and can be used to help infer the locomotion of fossil mammals of all sizes.

Despite the considerable research focus on identifying links between limb morphology and locomotion, historically few studies on the morphology of climbing in mammals account for the influence of phylogenetic relatedness and intraspecific variation, and the study taxa are often limited to mammalian subclades. Thus, our aim is to robustly examine limb correlates of arboreality using advanced comparative methods and a phylogenetically broad mammalian dataset.

Here, we have three objectives. First, we briefly review the fossil record of arboreality from non-mammalian synapsids to early therian mammals (Mesozoic through early Cenozoic). Second, we use multilevel regression models to examine which limb measurements are the strongest predictors of arboreality in a large sample of extant mammals, using linear measurements that capture functional traits like in-lever/out-lever lengths and are easily obtained from fossil specimens. The results provide guidance for inferring climbing in extant and fossil taxa, and we provide suggestions for future studies that apply similar methods. Third, we apply the measurements and regression models to three fossil case studies of early mammalian clades that are inferred to have been mostly arboreal: Mesozoic Theria, Multituberculata, and Plesiadapiformes.

A central finding is that extant (and presumably fossil) arborealists rarely exhibit a full suite of limb adaptations for arboreality; rather, some of their traits appear to be adapted for arboreality whereas other traits are more similar to those of ground-dwelling terrestrialists. This highlights the difficulty of interpreting small-mammal morphology and emphasizes the importance of using multiple traits when inferring locomotor modes in fossil taxa. Further, we support previous findings that measures of digit length and elbow traits are especially powerful predictors of arboreality (e.g., Bloch and Boyer, 2002; Kirk et al., 2008; Chen and Wilson, 2015; Meng et al., 2017; Nations et al., 2019). Finally, our analyses of fossil taxa provide novel insight into the locomotor evolution of several extinct groups and cast doubt on the hypothesized prevalence of arboreality among early mammalian lineages.

## 2 Evolution of Arboreality in Early Mammals

### 2.1 Arboreality in Non-Mammalian Synapsids

Mammals are the crown members of a more inclusive clade, Synapsida, that evolved ca. 315 Ma (Kemp, 2005; Hellert et al. 2023). Non-mammalian synapsids, such as ‘pelycosaurs,’ ‘therapsids,’ and ‘cynodonts’ are inferred to have been primarily ground-dwellers (as was likely the case with most early amniotes; e.g., Sumida and Modesto, 2001) with sprawling postures and ancestral ‘reptile-like’ limb morphologies (e.g., Romer, 1922; Romer et al., 1940; Jenkins, 1971; Kemp, 2005; Lungmus and Angielczyk, 2019). Although arboreality has been inferred in a few early synapsid taxa, such as the varanopid ‘pelycosaur’ *Ascendonanus* (Spindler et al., 2018) and the anomodont *Suminia* (Fröbisch and Reisz 2009, Fröbisch and Reisz 2011), early synapsids seem to have been primarily ground-dwellers.

Compared to earlier synapsids, there appears to have been a slightly increased prevalence of climbing adaptations among early mammaliamorphans (early mammaliaforms and close relatives; e.g., Guignard et al., 2019; Chen and Wilson, 2015), which may have attended other ecomorphological evolutionary trends that occurred near the origin of Mammaliamorpha, including an overall reduction in body size (Hellert et al. 2023). For instance, Guignard et al. (2019) inferred scansorial adaptations in the brasilodontid mammaliamorphan *Brasilodon*. Nonetheless, evidence for strict arboreality (rather than simply a capacity for climbing) is largely absent among early mammaliamorphans, and some of the most evolutionarily successful early mammaliamorphans, the Tritylodontidae, are inferred to have been mostly semi-fossorial (e.g., Sues and Jenkins, 2006; Mao et al., 2021). Indeed, even among the early mammaliaforms, most of their locomotor strategies have been inferred to be relatively generalized, lacking in obvious adaptations for tree-dwelling but not precluding the possibility for climbing, especially among Morganucodonta (e.g., Jenkins and Parrington, 1976). The clearest instance of specialized arboreal adaptations among non-mammalian mammaliaforms is from the holotype of the docodontan *Agilodocodon scansorius*, which exhibits a gracile humerus, elongated phalanges, and mesiodistally short but anteroposteriorly tall distal phalanges indicative of dedicated climbing (Meng et al., 2015). All other docodontans with known postcranial skeletal elements, however, are much more robustly built and have been inferred to be terrestrial to semifossorial (Martin 2005; Panciroli et al. 2022), fossorial (Luo et al., 2015), or even semiaquatic (Ji et al., 2006). Thus, based on the available fossil evidence, it is unlikely that arboreal locomotion was an important factor in the early evolution of non-mammalian synapsids.

‘Haramiyidans’ are one potential exception to the ground-dwelling prevalence among non-mammalian mammaliaforms. The phylogenetic placement of ‘haramiyidans’ is highly contentious (Fig. 1), with some researchers suspecting they (or at least some members of the group, namely Euharamiyida) bear a close evolutionary relationship to the Multituberculata (a group generally considered to belong in crown Mammalia, which will be discussed below) (e.g., Bi et al., 2014; Han et al., 2017), whereas others consider them to be phylogenetically stemward of the mammalian crown (e.g., Luo et al., 2017; Huttenlocker et al., 2018) or polyphyletic with both stem and crown mammal lineages (Hoffmann et al., 2020A; King and Beck, 2020; Krause et al., 2020). We do not wade into that debate here, but if ‘haramiyidans’ (or more specifically, Eleutherodontidae) are indeed non-mammalian mammaliaforms then they would represent the clearest example of widespread arboreal adaptations (e.g., Meng, 2014) among a clade of non-mammalian synapsids. Known primarily from isolated teeth in most places in the world, exceptionally preserved ‘haramiyidan’ skeletons from the Jurassic Yanliao Biota of China (ca. 160 Ma) have revealed pronounced arboreal adaptations among seven species of eleutherodontids (Bi et al., 2014; Luo et al., 2017; Meng et al., 2017; Han et al., 2017; Grossnickle et al., 2020), including at least three genera exhibiting gliding adaptation (including the soft-tissue preservation of the patagium, or ‘wing membrane’) (e.g., Luo et al. 2017; Meng et al., 2017; Han et al., 2017; Grossnickle et al., 2020) and phalangeal dimensions suggesting roosting behavior (Meng et al., 2017). Indeed, all but one (*Megaconus mammaliaformis*, which some contend is a multituberculate; e.g., Meng et al., 2014) known ‘haramiyidan’ skeleton discovered to date exhibit what have been interpreted to be adaptations for tree-dwelling (Zhou et al., 2013).

**Figure 1:**
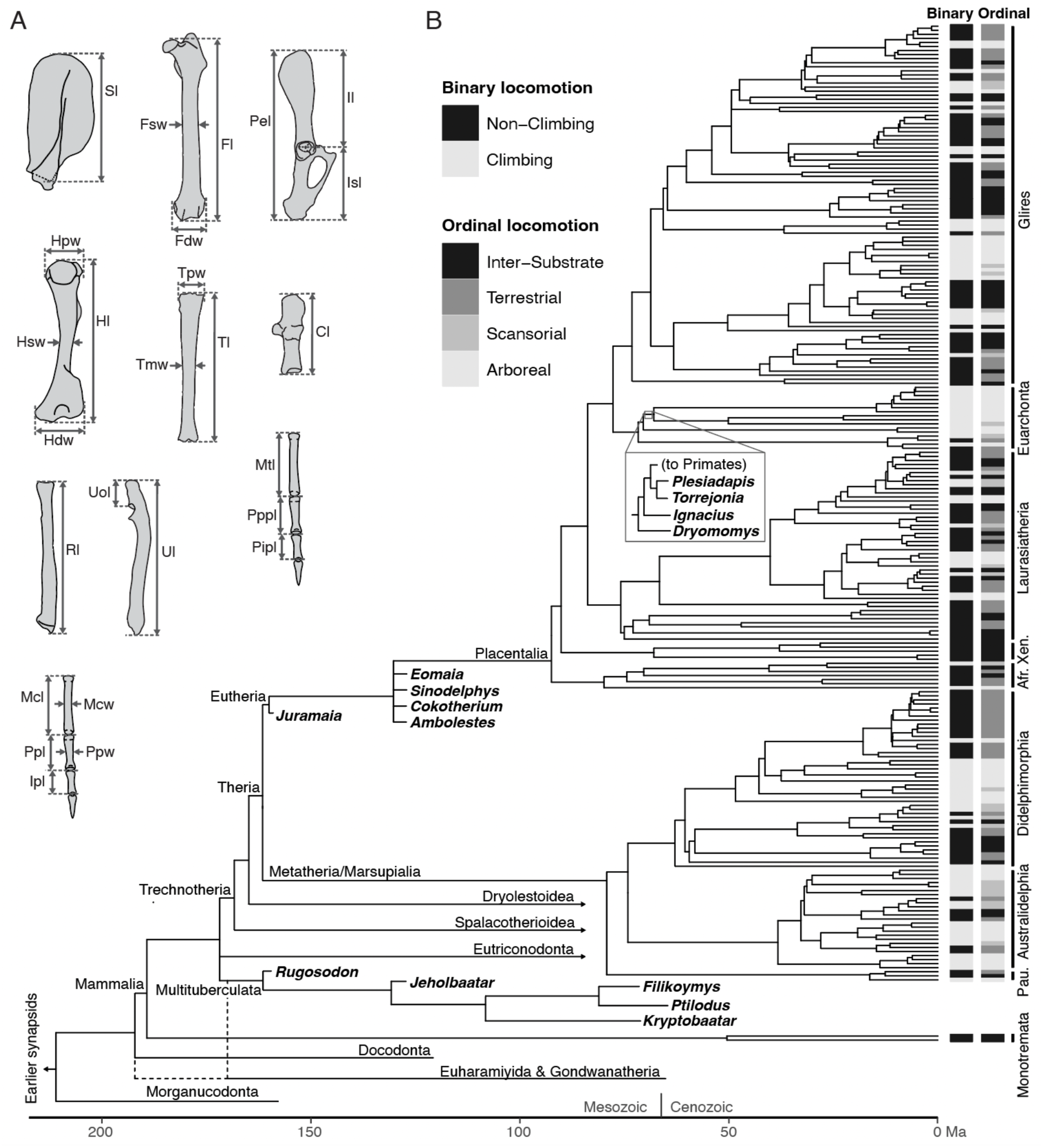
*A*, Linear measurements of the mammalian appendicular skeleton collected for this study. The illustrations are modified from those in Chen and Wilson (2015). *B*, Phylogeny of extant taxa (Upham et al., 2019) with extinct lineages grafted onto the tree. Fossil genera analyzed in this study are in bold. The topology of extinct lineages is based on several sources, including Bloch et al. (2007), Huttenlocker et al. (2018), Mao et al. (2019), Krause et al. (2020), and Weaver et al. (2021). The heatmaps show the binary (left) and ordinal (right) locomotor mode of each extant taxon in our dataset. Mammal clades are labeled at the ordinal, superordinal or grandordinal level. Pau. = Paucituberculata, Afr. = Afrotheria, Xen. = Xenarthra. Measurement abbreviations: Sl = Scapula Length, Fl = Femur length, Fsw = Transverse femur diameter, Fdw = Femur distal end width, Pel = Pelvic length, Il = Illium length, Isl = Ischium length, Hl = Humerus Length, Hpw = Humerus proximal end width, Hsw = Transverse humerus diameter, Hdw = Humerus distal end width, Tl = Tibia length, Tmw = Transverse tibia diameter, Tpw = Tibia proximal end width, Cl = Calcaneal length, Rl = Radius length, Ul = Ulna length, Uol = Olecranon process length, Mtl = Metatarsal III length, Pppl = Proximal phalanx III length pes, Pipl = Intermediate phalanx III length pes, Mcl = Metacarpal III length, Ppl = Proximal phalanx III length, Ipl = Intermediate phalanx III length, Mcw = Metacarpal III width, Ppw = Proximal phalanx III width.

### 2.2 Arboreality in Non-Therian Mammals

The prevalence of climbing adaptations in fossil lineages increases in early crown mammals (i.e., non-therian mammals). Two major groups in particular, Eutriconodonta and Multituberculata, both exhibit a range of locomotor modes among their members (e.g., Chen and Wilson, 2015; Grossnickle et al. 2019), including multiple taxa inferred to be scansorial or arboreal. Among the eutriconodontans, likely tree-dwellers include *Volaticotherium antiquum* (a possible glider; Meng et al., 2006, but see Meng et al., 2017 and Grossnickle et al., 2020) and *Jeholodens jenkinsi* (inferred to be arboreal; Chen and Wilson, 2015). In contrast, other eutriconodontans have been reconstructed as terrestrial and semifossorial (e.g., Jenkins and Schaff, 1988; Martin et al., 2015, Chen and Wilson, 2015), strongly suggesting that Eutriconodonta achieved considerable locomotor diversity.

Compared to other groups of early crown mammals, the locomotor habits of multituberculates have been explored in detail. Mostly based on postcranial specimens from the Paleocene of North America, multituberculates have historically been interpreted as being scansorial or arboreal (Simpson, 1926; Simpson and Elftman, 1928; Dieschl, 1964; Jenkins and Krause, 1983; Krause and Jenkins, 1983; Rowe and Greenwald, 1987), although the earliest interpretation of multituberculate locomotion suggested they were saltatorial (Gidley, 1909; a hypothesis later disputed by Simpson [1926]). More recent evaluations of multituberculate specimens, however, have suggested a greater diversity of locomotor modes. The oldest multituberculate represented by a postcranial skeleton (*Rugosodon*; Late Jurassic, ca. 160 Ma) was interpreted as terrestrial (Yuan et al., 2013; Luo et al., 2016), but it was later inferred to have been scansorial (Chen and Wilson, 2015). Most Cretaceous taxa have also been interpreted to be terrestrial, with some exhibiting burrowing adaptations (*Filikomys*; Weaver et al., 2021) and others exhibiting leaping adaptations (*Kryptobaatar*, *Nemegtbaatar*; Kielan-Jaworowska and Gambaryan, 1994). Only one Cretaceous taxon has been interpreted to be truly arboreal (*Sinobaatar*; Chen and Wilson, 2015). Committed fossoriality has also been inferred for one multituberculate, ?*Lambdopsalis bulla*, from the early Paleogene of China (Kielan-Jaworowska and Qi, 1990), and has been suggested for *Taeniolabis* from the Paleocene of North America (Simpson and Elftman, 1928). Most multituberculates represented by more fragmentary postcranial remains (i.e., femora and humeri) from the latest Cretaceous and earliest Paleocene of Montana have been inferred to be mostly arboreal with some terrestrial taxa interspersed (DeBey and Wilson, 2014; DeBey and Wilson 2017). In sum, multituberculates apparently exhibited a wide diversity of locomotor modes but may have more prevalently committed to arboreality after the Cretaceous-Paleogene mass extinction.

The locomotor habits of Gondwanatheria, an enigmatic group from the southern hemisphere that may be closely related to multituberculates (Krause et al., 2020; Hoffmann et al., 2020A) or eleutherodontid ‘haramiyidans’ (e.g., Huttenlocker et al., 2018), are very poorly known because only one gondwanatherian skeleton has been discovered (Krause et al., 2020). Nonetheless, that specimen is robust in postcranial morphology and resembles ambulatory and semifossorial mammals (Hoffmann et al., 2020B).

Close relatives of therians (i.e., non-therian Trechnotheria) such as spalacotheroids and dryolestoids are similar to multituberculates in that they appear to have exhibited a diversity of locomotor strategies, but evidence for individuals with committed or obligate arboreal locomotion are rare. The Spalacotheroidea (‘symmetrodonts’ with acute-angled molar cusps) include hypothesized semifossorial (*Akidolestes cifellii*; Chen and Luo 2013), terrestrial (*Maotherium*; Rougier et al. 2003), and scansorial (*Zhangheotherium quinquecuspidens*; Chen and Wilson 2015) representatives. Postcranial specimens of non-therian cladotherians are very rare, but *Henkelotherium* has been inferred to have been arboreal (e.g., Jäger et al. 2020),*Dryolestes* is likely to have been terrestrial (Jäger et al. 2020), and *Necrolestes* was inferred to have been fossorial (Rougier et al., 2012).

### 2.3 Arboreality in Mesozoic Theria

Arboreality has featured prominently in discussions about the evolution of therian mammals, especially marsupials and placentals (Huxley, 1880; Dollo, 1899; Bensley, 1903; Matthew, 1904; Haines, 1958; Szalay, 1981; Szalay and Sargis, 2001; Luo et al., 2003; O’Leary et al., 2013; Hughes et al., 2021). Among the earliest representatives of the more inclusive marsupial and placental lineages—Metatheria and Eutheria, respectively—their postcranial skeletons have been interpreted to exhibit arboreal or, at least, scansorial adaptations (Ji et al., 2002; Luo et al., 2003; Luo et al., 2011; Chen and Wilson, 2015; Meng et al., 2017; Bi et al., 2018).

Consequently, arboreality has been inferred to be the plesiomorphic condition among therian mammals (Ji et al., 2002; Luo et al., 2003; Luo et al., 2011). However, a split in locomotor specialization has been proposed between Metatheria and Eutheria, with eutherians exhibiting more terrestrial adaptations and metatherians exhibiting more arboreal adaptations (Huxley, 1880; Bensley, 1903; Dollo, 1899; Haines, 1958; Szalay, 1981; Szalay, 1984; Argot, 2001; Argot, 2002; Williamson et al., 2014). Though many of these arguments are based on early Cenozoic to Recent specimens, more fragmentary Cretaceous remains have been used to argue for an ancestral, arboreal locomotor mode in Theria (e.g., Godinot and Prasad, 1994; Prasad and Godinot, 1994; Goswami et al., 2011; Chester et al., 2012). Thus, although there is certainly evidence for terrestrial adaptations among Cretaceous and early Paleocene eutherians (e.g., Szalay and Decker, 1974; Kielan-Jaworowska, 1978), the current consensus is that climbing was prevalent in the earliest evolutionary history of Theria (e.g., (Ji et al., 2002; Luo et al., 2003; Luo et al., 2011; Bi et al., 2018).

### 2.4 Arboreality in early Paleogene Theria

Given the fragmentary nature of latest Cretaceous and earliest Paleocene postcranial remains, it is difficult to parse with confidence trends in therian locomotor trends across the Cretaceous-Paleogene boundary, though some studies have utilized fragmentary remains to infer patterns in morphological disparity that hint at some selection for locomotor mode (DeBey and Wilson, 2014; DeBey and Wilson, 2017). Based on inferred ancestral states of extant mammals and trait transition rates, it has been proposed that there was an ‘arboreality bottleneck’ at the K-Pg boundary associated with the fallout from the Chicxulub bolide impact (Hughes et al., 2021; and see Wu et al. [2017] for evidence of a post-K-Pg bottleneck for multiple ecological traits including arboreality); however, inferred arboreal lineages like the Plesiadapiformes (stem primates) likely arose in the Cretaceous (Wilson Mantilla et al., 2021) and they persisted into the Paleocene, so the mechanisms proposed for the ‘arboreality bottleneck’ still require investigation. In any case, the earliest Paleocene mammals for which there are records, postcranial skeletons from the San Juan Basin of New Mexico, are universally considered to have exhibited robust body plans, implying minimal agility and a more ambulatory, terrestrial mode of locomotion (Matthew, 1937; Rose, 2006; Shelley et al., 2021). Nonetheless, in South America, some early Paleocene metatherians have been interpreted to be scansorial (e.g., Argot, 2001; Argot, 2002), and it is possible that widespread terrestriality was merely a feature of the earliest archaic ungulates (i.e., ‘condylarths’; Archibald, 1998; Shelley et al., 2021), which dominate the postcranial fossil record of the San Juan Basin (e.g., Matthew, 1937), rather than early Paleocene mammals more broadly.

Among eutherians, the clear exception to these more terrestrial early Paleocene North American taxa are the Plesiadapiformes, among which even their earliest representatives exhibit distinct arboreal adaptations (Chester et al., 2015; Chester et al., 2019). As the Paleocene progressed, however, the prevalence of arboreal locomotion among eutherians increased and perhaps became the most common locomotor strategy (e.g., Macleod and Rose, 1993; Argot, 2013), revealed by more complete postcranial specimens in the middle–late Paleocene and early Eocene of North America (e.g., Rose, 2006). This increased prevalence of arboreality likely tracks the emergence of dense, closed-canopy, angiosperm-dominated forests in the early Paleogene (e.g., Carvalho et al., 2021; Benton et al., 2022).

In summary, non-mammalian synapsids and non-therian mammals mostly exhibited terrestrial or semifossorial locomotor strategies, with ‘haramiyidans’ and Paleocene multituberculates being exceptional in their prevalence of arboreal adaptations. In contrast, early therians are viewed as ancestrally arboreal, and indeed the earliest therian postcranial skeletons have been interpreted as exhibiting adaptations for climbing. Despite arguments that extant placentals derived from a terrestrial ancestor and extant marsupials from an arboreal ancestor, arboreality has nonetheless become synonymous with the locomotor preferences of early therians. At the K-Pg boundary, there may have been an arboreal bottleneck, through which mostly terrestrially adapted taxa survived, and this scenario is supported by the more robustly built early Paleocene archaic ungulates. However, the scansorial earliest Paleocene metatherians and the strictly arboreal plesiadapiforms, which likely arose in the Late Cretaceous, cast doubt on this evolutionary scenario. Nevertheless, by the middle–late Paleocene, arboreality likely became prevalent among mammalian communities, and this shift from more common terrestriality to arboreality likely tracked the emergence of diverse, angiosperm-dominated forest in the early Paleogene.

## 3 Predicting Climbing in Mammals

A central aim of this study is to examine which linear limb traits are the strongest predictors of arboreality in mammals to better inform our locomotor predictions of extinct species. To do this, we use a large sample of extant mammalian species with available skeletal material and known locomotor affinities to identify informative limb measurements. In this section, we explain the sampling, measurements, and predictive modeling of the extant species. In the next section (4), we use the measurements with strongest predictive potential to assign climbing probabilities to a selection of 14 mammalian fossil taxa from three mammalian lineages that have featured prominently in discussions about the evolution of arboreality in the Mesozoic and early Cenozoic (Fig. 1).

### 3.1 Sampling of Morphological Measurements from Extant and Extinct Mammals

We compiled 49 linear measurements of the appendicular skeleton from 427 specimens of 236 extant mammalian species (including the recently extinct thylacine). The samples stem from 70 families in 21 orders of mammals and include two species from two families of monotremes, 72 species from 15 families in eight orders of marsupials, and 163 species from 53 families in 13 placental orders. The primary sources for measurements are Chen and Wilson (2015), Meng et al. (2017), Nations et al. (2019), Grossnickle et al. (2020), Weaver and Grossnickle (2020), and Pevsner et al. (2022). Of the 49 measurements, 12 are complete for all 427 specimens, and 14 have less than 50% missing data. We excluded measurements that had more than 50% missing data, resulting in a final set of 26 measurements used in our analyses. The measurements are illustrated and described in Figure 1 (full list available in Table S1). Because we focus on arboreal adaptations and Mesozoic–early Cenozoic mammals (which, to date, do not appear to have been volant or fully aquatic), we included only terrestrial and amphibious, non-volant mammals (we excluded bats, seals, whales, dugongs, etc.). When available, we recorded the mass in grams from museum tags or field notes for measured specimens. We supplemented the mass data with estimates from panTheria (Jones et al., 2009) and, in a few cases, from EltonTraits (Wilman et al., 2014) or additional publications (Table S1). At the extremes of the locomotor spectrum, sampled species include fully arboreal primates, amphibious muskrats and otters, and burrowing moles and gophers; however, most specimens do not display these sorts of extreme adaptations to select locomotor modes. Ultimately, this large dataset provides ample morphological, ecological, and phylogenetic diversity to address a range of questions regarding locomotor adaptations in mammals.

Because we aim to predict locomotion in extinct mammals, our dataset is tailored to maximize the applicability to fossil taxa. Our extant sampling consists primarily of small-to-medium-sized species as these are most appropriate as analogs of early mammal fossil taxa, which are relatively small (e.g., Chen and Wilson, 2015). The body mass of our sample includes three species over 100 kg and 34 above 5kg. The remaining 203 species, 86% of the sample, are at or under 5 kg, the cutoff that is often used to categorize “small mammals” (e.g., Merritt, 2010). Further, we use linear measurements rather than 3D measurements because the skeletons of many fossil taxa are compressed, flattened, or embedded in rock, making 3D analyses challenging to impossible. An additional benefit of using linear measurements is that they are commonly used in mammalian ecomorphological studies (Dunn, 2018), and this provides us the opportunity to test, in a systematic way, which measurements or types of measurements have the greatest predictive accuracy.

### 3.2 Describing Locomotion in Extant Species

To analyze which measurements best predict climbing, we must first separate the climbers from the non-climbers in our data set. Rather than using many discrete categorical locomotor modes (e.g., Samuels and Van Valkenburgh, 2008; Chen and Wilson, 2015; Dunn, 2018), we used two different classification schemes tailored to climbing behavior: (1) binary categories (‘climb’ and ‘no climb’) and (2) ordinal locomotor rankings, reflecting a gradient of climbing preference, or climbing probability (Inter-substrate, Terrestrial, Scansorial, and Arboreal). These two classification schemes require different modeling methods, which are discussed below. For the binary categorization scheme (‘climb’ and ‘no climb’), we assigned scansorial (see definition below), arboreal, and gliding taxa into the “climb” category. In essence, this places all the non-climbing taxa, and their disparate morphologies, into a single pool, and separates the climbing species (Weaver and Grossnickle, 2020). Our final count is 95 species (150 specimens) in the “climb” group, and 142 species (277 specimens) in the “no climb” group.

For the ranked ordinal classification scheme, we assigned each species to one of four ascending levels of arboreal locomotion, with each level representing increasing reliance on climbing: (1) Inter-substrate: The lowest rank, including taxa that actively forage and/or burrow and nest underground or in water, and are highly unlikely to climb; (2) Terrestrial: The second rank, comprising taxa that do not burrow deeply (e.g., nesting in leaf litter) and, while potentially capable, are not known to climb for food or shelter under normal circumstances; (3) Scansorial: The third rank, encompassing taxa that actively forage and/or nest on both the ground and in trees, demonstrating moderate climbing abilities; (4) Arboreal: The highest rank, including taxa that forage and nest primarily in trees, for which climbing is a critical aspect of their biology. The ranking method is similar to that from Nations et al. (2019). This ranked system reflects a gradual increase in arboreality and climbing dependence across the four levels, or “continuous probability categories,” rather than discrete, unordered categories. However, we use these four locomotor classification terms rather than a numeric ranking as they, and other similar locomotor classifications, are often used as discrete categories in other studies and are familiar to the field (e.g., Samuels and Van Valkenburgh, 2008; Chen and Wilson, 2015; Meng et al., 2017; Dunn 2018). Our final count is 65 inter-substrate species (100 specimens), 77 terrestrial species (177 specimens), 25 scansorial species (42 specimens), and 70 arboreal species (108 specimens). Sources for these locomotor rankings are based on previous quantitative studies on locomotor modes (e.g., Samuels and Van Valkenburgh, 2008; Chen and Wilson, 2015; Nations et al., 2019; Grossnickle et al., 2020; Weaver and Grossnickle, 2020), additional primary literature, Nowak (1999), and online databases (e.g., the Animal Diversity Web, www.animaldiversityweb.org, and sources within). All sources are provided in Table S1.

Although our ordinal ranking method differs from traditional categorical classification, we acknowledge that any categorizing taxa is overly simplistic—no mammal engages in just a single behavior. Even taxa that are specialized for specific locomotor behaviors may occasionally exhibit other behaviors, such as tree squirrels occasionally running, digging, or swimming (e.g., Nice et al., 1956; Hauser, 1964). This can complicate analyses because (1) many taxa may have adaptations for performing multiple locomotor modes (i.e., generalist morphologies), making it challenging to identify form-function relationships across taxa, and (2) it can lead to misclassification of locomotor modes because various studies could report different behaviors for any given species. We help to address this issue by using two different methods (binary and ordinal ranks); differences in results of these analyses would suggest that results are sensitive to classification and ranking schemes. Additionally, focusing our locomotor categories on climbing alone (a binary climb/no-climb and an ordinal ranking of climbing probability) rather than locomotion more generally, we can identify traits that covary with climbing rather than investigating morphological differences among many discrete locomotor modes, as is common in our field. Further, we emphasize throughout our study that results improve with the inclusion of a greater number of morphological traits in analyses. Thus, even if classification-related issues disrupt some form-function signals in our study, we believe that this can be overcome by using large samples of taxa and multiple morphological measurements.

Scansorialists are especially challenging taxa to classify because they exhibit both ground-dwelling and tree-dwelling behavior. Here, we define scansorialists as taxa that actively forage and/or nest on both the ground and in trees. This definition is admittedly unsatisfactory because climbing versus non-climbing behavior is a spectrum; the amount of climbing behavior needed to constitute scansoriality is subjective, and limited or contradictory natural history information for many species further magnifies the issue. However, we doubt that these problems with scansorial classifications significantly influence our results and conclusions. For instance, taxa that we define as scansorialists only comprise ∼10% of our dataset (25 of 237 species, 42 of 427 specimens); thus, misclassifications of some scansorialists should not significantly influences statistical results. Further, misclassifications may be working against our results, which tend to show strong links between morphology and locomotion, and future studies that update and improve our classifications may observe even stronger limb-locomotion relationships than those found here.

### 3.3 Calculating Morphological Metrics

A linear measurement from the skeleton of a mammal cannot be directly interpreted without knowledge of the individual’s size. The need to condition on body size has led to a panoply of methods (e.g., Mosimann, 1970; Jungers et al., 1995; Klingenberg, 2016), some contentious (see Freckleton, 2009), that minimize the influence of body size, ranging from calculating ‘size-free’ functional indices (i.e., ratios of two or more measurements; Hildebrand, 1985a; Hildebrand, 1985b; Elissamburu and Vizcaíno, 2004; Sargis, 2002a; Samuels and Van Valkenburgh, 2008; Chen and Wilson, 2015; Dunn, 2018; Woodman, 2023) to using estimates of residuals from a univariate body-size regression as input data in a secondary analysis (e.g., Kilbourne, 2017).

Here we focus on three means of conditioning our inferences on size: (1) regression models of log-transformed linear traits with log-transformed body size (either body mass or geometric mean) as a second predictor variable (hereafter ’linear metrics’), (2) skeletal ratios (i.e., functional indices) of two or more measurements, and (3) log-shape ratios (LSRs) with the geometric mean of multiple measurements as the denominator (Mosimann, 1970; Claude, 2013). We chose these methods based on statistical first principles, available data, and commonly used methods in the literature. For the first method, we natural log-transformed both the linear measurements and mass to place them on the same order of magnitude. Body size can influence both the length of the phenotypic measurement and the locomotion of a species, representing a ‘fork’ in the causal relationship that needs to be conditioned upon (Cinelli et al., 2022), and therefore we built regression models (details below) using both log-transformed linear measurements and, for linear metrics, log-transformed mass and, in a separate set of models, log-transformed geometric mean as predictor variables.

From our 26 linear measurements, we calculated 17 functional indices (Table 1), all of which were used in previous studies (Lemelin, 1999; Sargis, 2002a; Elissamburu and Vizcaíno, 2004; Samuels and Van Valkenburgh, 2008; Hopkins and Davis 2009; Chen and Wilson, 2015; Woodman and Stabile, 2015; Dunn, 2018; Nations et al., 2019; Woodman, 2023). These indices are typically length-by-width or length-by-length type ratios thought to describe the proportions of skeletal elements and have been shown to minimize the correlation with body size (Weisbecker and Schmid 2007). For example, Femoral Robustness Index (FRI) is the midshaft width of the femur divided by the overall length of the femur. A species with a stout, robust femur will have a higher FRI than one with a long, gracile femur, regardless of body size. Each functional index was used as a predictor variable in the regression models (see Table 1 for a complete list of indices and their functional significance for climbing).

**Table 1:**
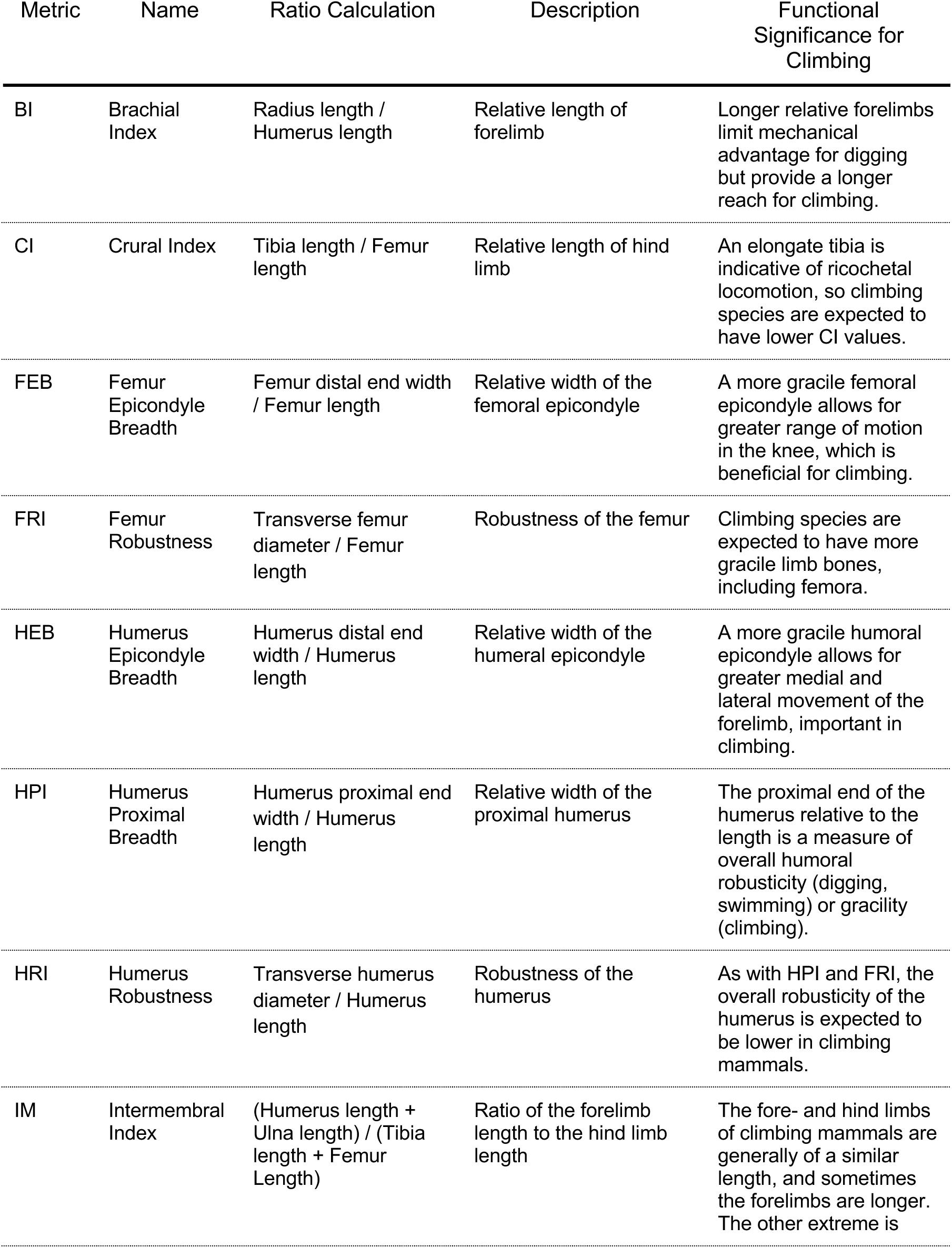
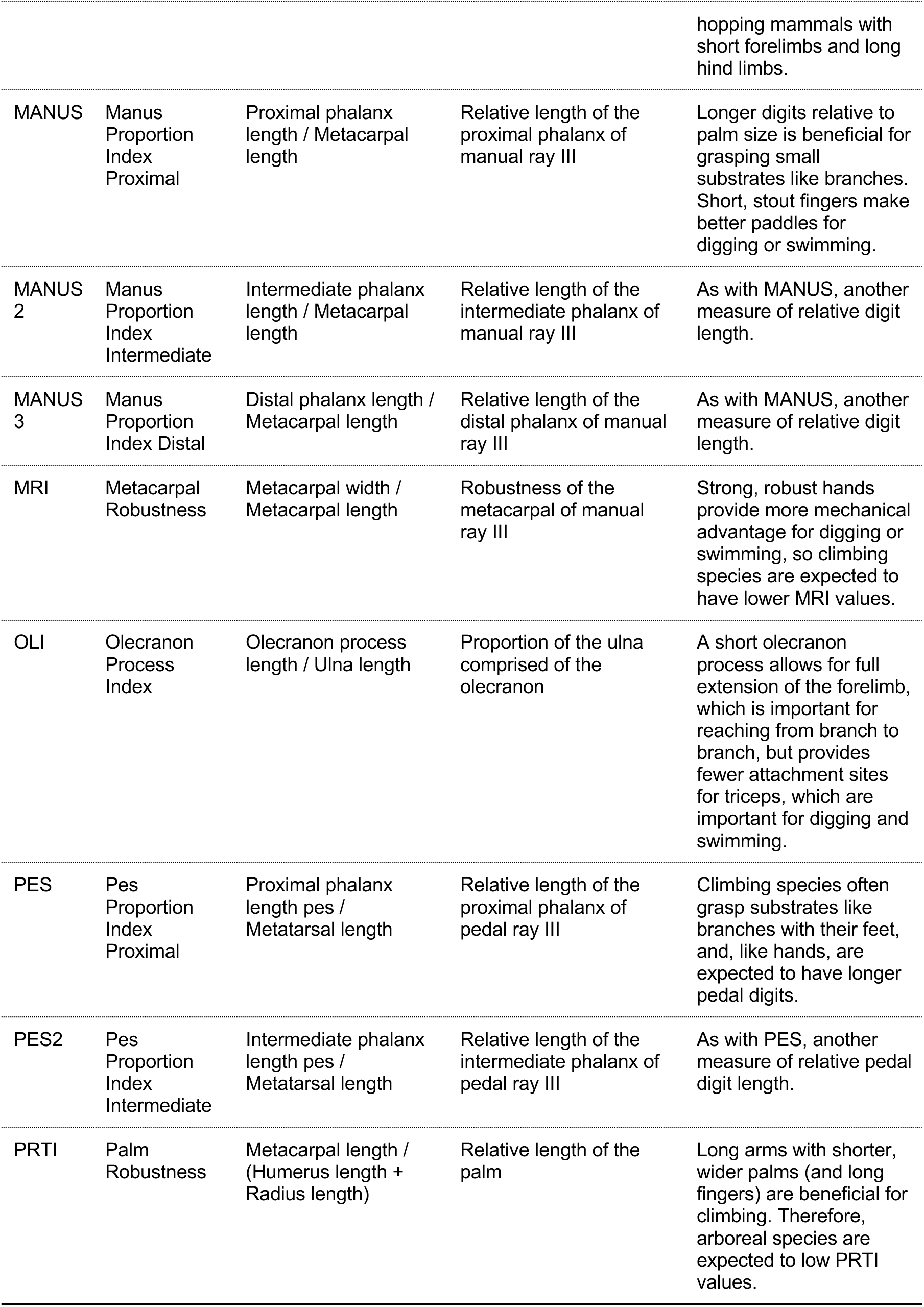
Names, calculations, descriptions, and functional significance of the 17 Functional Indices examined in this study. Citations describing the functional significance of each index are found in section 3.3.

To calculate LSRs, we first calculated the geometric mean for each specimen by multiplying the values of 12 measurements together (Femur distal end width, Femur length, Femur midshaft width, Humerus distal end width, Humerus length, Humerus midshaft width, Olecranon process length, Radius length, Scapula length, Tibia length, Tibia midshaft width, Ulna length), then taking the 12th root of the value. We chose the 12 measurements because they had little (<3 species) to no missing data across our extant dataset. All these measurements are from the appendicular skeleton, making our geometric mean values a proxy of limb and girdle size rather than overall body size. The geometric mean of measured skeletal elements is a commonly used proxy for size (e.g., Claude, 2013; Kilbourne, 2017; Price et al., 2019, Weaver and Grossnickle, 2020; Pevnser et al., 2022) that can be especially useful when ample skeletal material is available but reasonable body mass measurements are not (e.g., fossil specimens; Grossnickle et al., 2020). We calculated LSRs of each of the 26 linear measurements by dividing each linear measurement value by the geometric mean of the specimen, then log transforming the product. This method of removing size preserves any allometry that is present in the raw data (Mosimann, 1970; Junger et al., 1995; Klingenberg, 2016) and has been used in numerous studies on mammals and other vertebrates (Claude, 2013; Price et al., 2019; Weaver and Grossnickle, 2020; Pevsner et al., 2022).

In total, we calculated 69 skeletal metrics (26 log-transformed linear traits, 17 functional indices, and 26 LSRs) and two body size metrics (log-transformed mass and geometric mean) that are examined in subsequent analyses. Moving forward, we use initial capitalization for linear metrics (e.g. Ppl), all caps for indices (e.g. OLI), and label log-shape ratios with the prefix LSR.

### 3.4 Modeling Relationships Between Skeletal Elements and Locomotion

To determine which skeletal limb metrics best predict climbing in mammals, and then to use these metrics to estimate the locomotor mode of extinct taxa, we use probabilistic regression modeling, where the effect sizes of each measurement on climbing can be generated and model accuracy can be evaluated through predictive sampling of the training data.

#### 3.4.1 Constructing Bayesian Models

Here we use two different regression families: logistic regression for the binary climbing categories, and ordinal regression for our ranked locomotor descriptions. Logistic regression models, henceforth “binary models,” estimate the effect of the skeletal metric on climbing, and can generate a predictive distribution of climbing probability, on the 0–1 scale, for each species in the dataset or for data that lack locomotor information, like fossils. Cumulative ordinal regression models assume the response variable (locomotion) is drawn from a continuous distribution split by thresholds. Our ordinal models used the four ordered locomotor rankings of Inter-Substrate, Terrestrial, Scansorial, and Arboreal as described above, and therefore estimated three thresholds, one between each of the four ranks. Ordinal models estimate the effect of the skeletal metric on climbing and generate a predictive distribution of the probability of belonging in each rank for each species, which can be used to predict the locomotor rank of species that are not in the original dataset, such as fossils.

For both binary and ordinal regression families, we constructed multilevel models that include the locomotor value as the response variable, and the phenotypic metric as the population-level predictor (linear metric models also included log-transformed mass or geometric mean as a second predictor). Because many species are represented by multiple specimens, we include the species as a group-level effect, which uses partial pooling to estimate a mean value for each species, then takes these partially pooled effects to estimate the overall effect of the measurement on the locomotor value. The resulting posterior distribution of species-level variance provides an estimate of the intraspecific variability in the metrics. We also include the phylogenetic correlation matrix, calculated from a maximum clade credibility (MCC) consensus phylogeny generated from 1000 randomly chosen trees from Upham et al. (2019), as an additional group-level term. We used the multilevel-model approach to phylogenetic regression, outlined in detail in de Villemereuil and Nakagawa (2014), which jointly estimates the influence of the phylogeny on the distribution of the predictors, and conditions on the strength of that correlation. Like the species group-level effect, the phylogenetic group-level effect generates a variance estimate that indicates the contribution of the phylogenetic correlation matrix on the morphological metrics.

Of our 69 metrics (26 linear metrics, 17 functional indices, and 26 LSRs), 36 have some missing data (Table S1). Rather than dropping numerous samples from each of the models with missing data, we leveraged the power of Bayesian modeling to jointly estimate the missing data within the larger binary or ordinal modeling framework. This method requires jointly estimating two linked models within a single, multivariate model. The first is a Gaussian linear regression that, for each model iteration, estimates the value of each missing measurement for each sample using 13 predictors—12 linear measurements with no missing data (the 12 used in the GM calculation, see above) and body size as log-transformed mass—along with the phylogenetic placement of the individual as predictor variables. For each iteration, the predicted missing values are added to the full dataset, and the binary or ordinal model with the same model structure and covariates that are described above estimates the effect of the metric on climbing (Bürkner 2018; McElreath, 2020). We used the mi() function in the *R* package *brms* v2.19.6 to build these missing data models (Bürkner, 2018). Together we ran models for each combination of response distribution and morphological metric. Before fitting the models, all variables were scaled to a mean of zero and a standard deviation of one. Regularizing priors were fit on all variables to discourage the searching of unrealistic parameter space. Models were fitted using functions in the *brms R* package, which uses the Bayesian probabilistic programming language Stan. Each model was run for four chains, with 2,000 iterations of warm-up and 2,000 iterations of sampling. Chain convergence was verified with the Gelman-Rubin *R̂* statistic (Gelman and Rubin, 1992). All model scripts and Supporting Information are available at https://github.com/jonnations/Locomotion_Book, and are archived at https://zenodo.org/doi/10.5281/zenodo.10144438.

#### 3.4.2 Binary Model Results

All binary model chains effectively converged, as indicated with an *R̂*≤ 1.001 for each parameter. Morphological metric effect sizes ranged substantially from robust effects for several manual and pedal measurements (Proximal phalanx length [Ppl], Pes Proportional Index Proximal [PES], Manus Proportional Index Proximal [MANUS]) and humeral, ulnar, and olecranon measurements, to effect sizes centered on zero with large variances (i.e., Femur midshaft width [Fmw], Tibia proximal end width [Tpw], etc. Fig. 2, S1, Table S2). Importantly, the effects highlight the directionality of the relationships; digits and long bone lengths, both absolute and relative, are positively related to climbing, and olecranon length and humerus robusticity are negatively related to climbing. In all models, the intra-specific variance parameter is larger than the phylogenetic variance parameter, indicating that within-species variation contributes more to the overall model variance than the phylogenetic correlation between traits.

**Figure 2:**
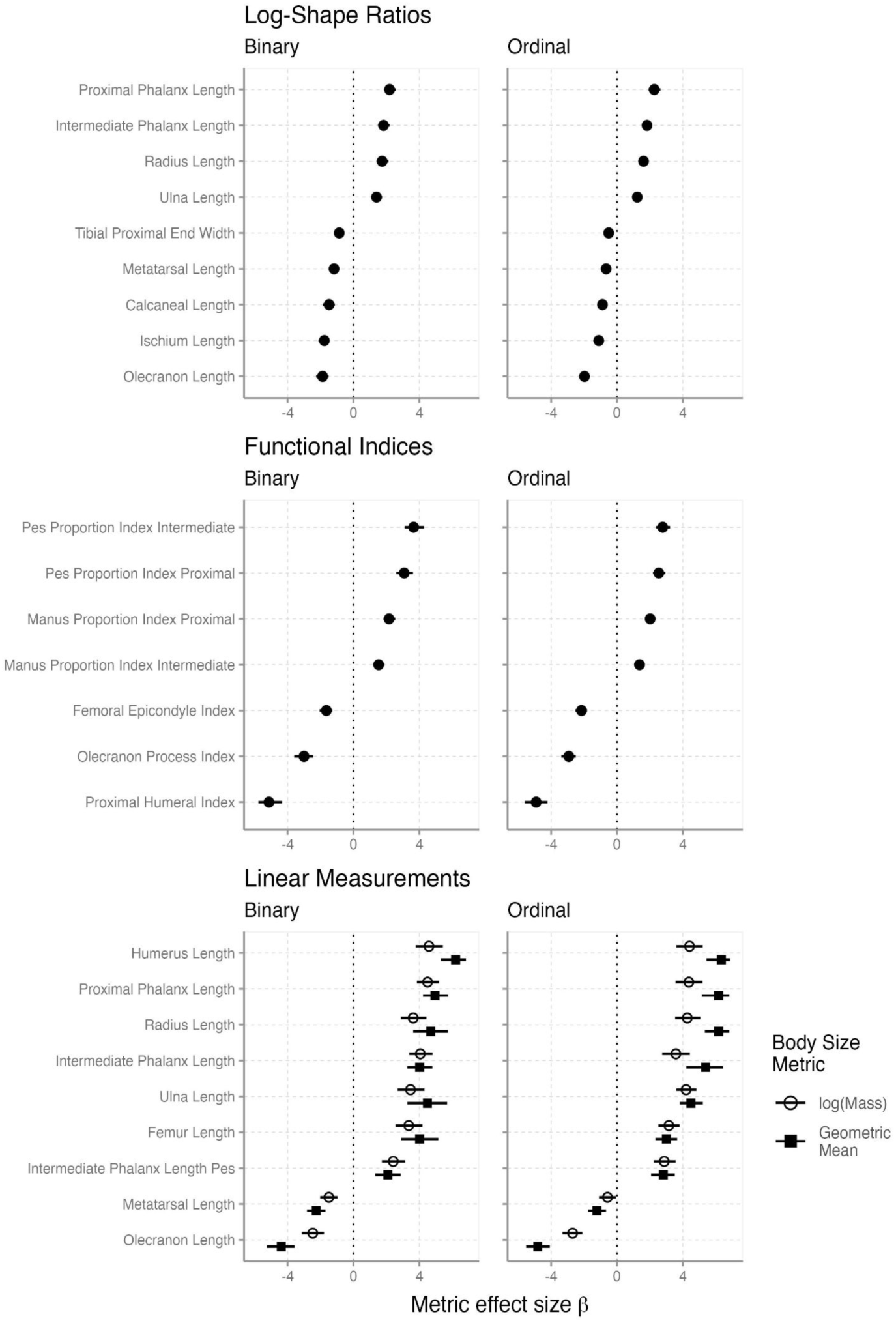
The effect sizes (log-odds scale) from the highest-accuracy phenotypic metrics from each of the three metric size-correction classes. The left column shows effect sizes from binary models, and the right column from ordinal models. Points show the median estimate, and interval lines show the 89% probability distribution. Positive effect sizes means a positive relationship with metric value and climbing, and a negative effect size indicates a negative relationship. Each of the metrics has a predictive accuracy ≥ 70% when generating predictions without phylogenetic information. Results from all models can be found in the Zenodo repository as “O_Effects_All.pdf” and “B_Effects_All.pdf”

#### 3.4.3 Ordinal Model Results

As with the binary models, all the ordinal model chains properly converged (*R̂*≤ 1.001). The morphological metric effect sizes from the ordinal models are very similar to those of the binary models, with manual and pedal metrics, and humeral, radial, and ulnar metrics having the largest effect sizes, both positive and negative (Fig. 2, S2, Table S3). A large effect size in ordinal regression generally means good separation on the phenotype axis among the different rankings, or in other words, locomotor ranking can be predicted from morphological metric value with greater precision than for metrics with near-zero effect sizes. As in the binary models, the intra-specific variance parameter typically has a larger value than the phylogenetic variance parameter, however both values are generally higher than the same parameters in the binary models.

### 3.5 Testing the Accuracy of Predictive Models

Although the initial results of the Bayesian models appear promising for some metrics, secondary measures were taken to ensure that accurate predictions can be made. This includes verifying that parameter estimates fall within reasonable values and that models are not overfit to the data. Additionally, we are interested in which of the morphological metrics are the most accurate predictors of climbing in mammals with both a known and unknown phylogenetic position. Once accurate metrics are identified, we construct binary and ordinal multiple regression models that include all the accurate predictors, and then compare their predictions to the models with single morphological predictor variables.

#### 3.5.1 Testing for Overfitting and Accuracy

We used two different but complementary methods to validate the predictive ability of our models. First, we tested for overfitting. Overfitting occurs when the model learns “too much” from the data, resulting in near-perfect predictions on the original data, but rendering it incapable of generating accurate predictions for any new data (McElreath, 2020). We tested for overfitting using Pareto-*k* importance sampling in the leave-one-out (LOO) cross validation algorithm (Vehtari et al., 2017). LOO uses Pareto-*k* importance sampling to simulate running the model *n*-sample times, leaving out a single data point on each iteration, and then assigns each model an estimated log-predictive density score conceptually like an Akaike Information Criterion (AIC) or Watanabe-AIC (WAIC) score. LOO also assigns a Pareto-*k* value to each sample (i.e., specimen) in the model which describes the influence of that sample on the model’s posterior distributions. Although estimated log-predictive density scores cannot be directly compared among models due to differences in sample size stemming from missing data, we can examine the Pareto-*k* value assigned to each specimen in each model. The larger the Pareto-*k*, the more influential the data point, and numerous samples with high Pareto-*k* (>0.7) indicates model overfitting. We found that all models contained < 5% of samples with Pareto-*k* samples over 0.7, and many models contained zero.

Given the phenotypic and phylogenetic scale of the dataset, most samples with Pareto-*k* scores over 0.7 are unsurprising. For example, echidnas, which split from other therians by at least 170 Ma and are morphologically distinctive, had high Pareto-*k* scores for several metrics. Other examples point to kangaroos and springhares as influential outliers in pedal measurements, unsurprising given their extremely long metatarsals compared to close relatives. Tree hyraxes, which are adept climbers but do not resemble any other climbing mammals, are also outliers in some metrics. Species on the extreme edges of the morphological distributions, such as moles and burrowing rodents like mole rats, both of which have specialized and robust humeri, often had relatively high Pareto-*k* scores for humerus metrics. Rather than removing these few outlying samples from the models, we decided to include them because they represent realistic evolutionary trajectories, often found in phylogenetically distant taxa, and offer worthwhile comparisons for a fossil dataset.

Second, we estimated the accuracy of our models by using our extant data as a testing set by running it back through each model to get a prediction of the binary or ordinal locomotor scores for each extant species. This allows us to calculate the percentage of predictions that are correct, providing a measure of accuracy. To further test the predictive accuracy, we estimated prediction accuracy scores both with and without the phylogenetic correlation matrix by removing the phylogeny from the predictive model and rerunning our data as a testing set. Our models use the phylogenetic correlation matrix as a group-level correlation structure. Although modeling the relationship between morphology and phylogenetic distance is critical to understanding the role of phenotype in locomotion, oftentimes new data such as fossils cannot be accurately placed in time-scaled phylogenies of extant taxa. Furthermore, the phylogenetic distance between extant taxa and stem-clades, such as multituberculates, can be so great that, were the fossils to be accurately placed in the time-calibrated phylogeny, the model may be unable to confidently make any prediction on locomotion. Additionally, the modeling software *brms* does not allow for generating predictions with new correlation matrices not used in the original model (i.e., a new tree with fossil tips used in predictions)^1^. For the binary models, we calculated accuracy as the number of predictions that match the input data. We calculated the same accuracy for ordinal models, and we also calculated a percent of scores that are within one classification ranking of the input data (for example, an arboreal species that was predicted to be scansorial).

When generating predictions using phenotypic metrics and the phylogenetic group-level term, the binary models range from 94–100% accuracy. For the ordinal models, all metrics have between 83–88% accuracy, and 98–100% accuracy within one classification rank (e.g. an Arboreal taxon predicted to be Scansorial). Although our LOO analyses did not suggest strong overfitting, these high accuracy scores, even for morphological metrics with effect sizes that are centered on zero, indicate that the phylogenetic placement of the species is playing an outsized role in these models. In our predictions that excluded the phylogeny, the highest accuracy was in functional indices, linear metrics, and LSRs that describe digit lengths (Tables 2, S4), with the highest accuracy of 85% in the pedal digit length ratio PES2. The other ratios of manual and pedal digit length, MANUS and PES, as well as the ratio of the relative width of the proximal portion of the humerus HPI, also have over 80% accuracy. Additional measurements with high to moderate accuracy include the olecranon length ratio, ulna length, olecranon length, LSRs of ulna and olecranon length, and humerus length. The accuracy scores track with the strength of the predictor effect sizes, or the slope term of the phenotypic measurement in our generalized linear models (Figs. 2, S1, S2). The most accurate measures in the ordinal models were quite similar, including the manual and pedal digit ratios, the olecranon length ratio, and the proximal humerus ratio (Tables 2, S4), all of which had >55% accuracy in predicting the correct locomotor rank, and typically >80% accuracy in predicting within one rank of the input data.

Many of the most accurate linear measurements maintained similar accuracy whether using log(mass) or geometric mean as the body size descriptor, which is encouraging for predictions of fossil taxa that lack all the skeletal elements necessary for calculating geometric mean, or that may have unreliable body mass estimations. The LSRs of calcaneus length, olecranon length, and manual proximal phalanx length also performed well (Tables 2, S4), but we excluded these measurements from our predictive analyses (below) as the fragmented nature of our fossil samples precluded calculating LSRs for those taxa. Due to missing data in the fossil samples, we also excluded linear metric models that use geometric mean as the body size predictor. We proceed with multiple regression and fossil prediction using 14 metrics: PES2, HPI, PES, MANUS, Ppl, Hl, OLI, Pipl, MANUS2, Rl, Ipl, Uol, Fl, and Ul.

#### 3.5.2 Multiple Regression Models Using High-Accuracy Metrics

Multiple regression, or regression models with multiple predictors of a single response, is a highly effective predictive tool widely used across all scientific disciplines. To test the utility of multiple regression in predicting climbing from morphological data, we build two additional models, one binary and one ordinal, that use the 14 most accurate metrics as predictor variables (PES2, HPI, PES, MANUS, Ppl, Hl, OLI, Pipl, MANUS2, Rl, Ipl, Uol, Fl, and Ul; see Tables 1 and 2). Apart from the additional predictor variables, the model structure, priors, and execution remained identical to the single metric models. The models demonstrated effective convergence and appropriate Pareto-*k* values for all samples and will be used in predictions below.

**Table 2:** Names and descriptions of the linear measurements, functional indices, and log-shape ratios with the highest predictive accuracy among all metrics. Of the 69 metrics used in this study, these 23 are the best predictors of climbing in our dataset of 427 specimens of 237 mammal species, with a predictive accuracy over 70% for the binary model when the phylogeny is excluded from the predictions (see Figure 2). Each of these metrics have a predictive accuracy ≥95% when the phylogeny is included. Table S4 for contains the complete accuracy scores. The 14 linear measurements and functional indices in bold have the highest accuracy, the lowest quantity of missing data, and are used in extant and fossil predictions below.

| Linear Measurement | Name | Description |  | Accuracy |
| --- | --- | --- | --- | --- |
| FI | Femur length | Length of the femur |  | 73% |
| HI | Humerus Length | Length of the humerus |  | 76% |
| Ipl | Intermediate phalanx length - Manus | Length of the intermediate phalanx of the 3<sup>rd</sup> manual ray |  | 74% |
| Pipl | Intermediate phalanx length - Pes | Length of the intermediate phalanx of the 3<sup>rd</sup> pedal ray |  | 70% |
| Mtl | Metatarsal length | Length of the metatarsus of the 3<sup>rd</sup> pedal ray |  | 72% |
| Uol | Olecranon length | Length of the olecranon process of the humerus |  | 72% |
| Ppl | Proximal phalanx length - Manus | Length of the proximal phalanx of the 3<sup>rd</sup> manual ray |  | 81% |
| RI | Radius length | Length of the radius |  | 74% |
| UI | Ulna length | Length of the ulna, including olecranon process |  | 71% |
| Functional Index | Name | Ratio Calculation | Description | Accuracy |
| HPI | Humerus Proximal Breadth | Humerus proximal end width / Humerus length | Relative width of the proximal humerus | 83% |
| MANUS2 | Manus Proportion | Intermediate phalanx | Relative length of the | 75% |

|  | Index Intermediate | length / Metacarpal length | intermediate phalanx of manual ray III |  |
| --- | --- | --- | --- | --- |
| **MANUS** | Manus Proportion Index Proximal | Proximal phalanx length / Metacarpal length | Relative length of the proximal phalanx of manual ray III | 82% |
| **PES2** | Pes Proportion Index Intermediate | Intermediate phalanx length pes / Metatarsal length | Relative length of the intermediate phalanx of pedal ray III | 86% |
| **PES** | Pes Proportion Index Proximal | Proximal phalanx length pes / Metatarsal length | Relative length of the proximal phalanx of pedal ray III | 83% |
| **OLI** | Olecranon Process Index | Olecranon process length / Ulna length | Proportion of the ulna comprised of the olecranon | 75% |
| **Log-Shape Ratio of Linear Measurement** | **Name** | **Description** | **Accuracy** |  |
| LSR(CI) | Calcaneus length | Log(Length of the calcaneus / Geometric mean) | 72% |  |
| LSR(Ipl) | Intermediate phalanx length - Manus | Log(Length of the inter. phalanx / Geometric mean) | 71% |  |
| LSR(IsI) | Ischium length | Log(Length of the ischium / Geometric mean) | 72% |  |
| LSR(Mtl) | Metatarsal length | Log(Length of the metatarsus / Geometric mean) | 70% |  |
| LSR(OI) | Olecranon length | Log(Length of the olecranon / Geometric mean) | 72% |  |
| LSR(Ppl) | Proximal phalanx length - Manus | Log(Length of the prox. phalanx / Geometric mean) | 78% |  |
| LSR(Tpw) | Width of the proximal tibia | Log(Width of the proximal tibia / Geometric mean) | 70% |  |
| LSR(UI) | Ulna length | Log(Length of the ulna / Geometric mean) | 72% |  |

#### 3.5.3 Predicting Climbing in Extant “Model” Taxa

To further investigate our climbing predictions, we plotted the predicted results from 10 different species from our dataset, five climbing and five non-climbing. For the climbing species, we selected Derby’s Woolly Opossum *Caluromys derbianus*, the Philippine Colugo *Cynocephalus volans*, the Northern Flying Squirrel *Glaucomys sabrinus*, Douglas’s Squirrel *Tamiasciurus douglasii*, and the Common Treeshrew *Tupaia glis*. These species represent a variety of climbing styles and morphologies, and most have played an outsized role as extant climbing analogs of fossil species (Jenkins, 1974; Sargis, 2002a; Sargis, 2002b; Meng et al., 2006; Sargis et al., 2007; Thorington et al., 1998). All species have an ordinal ranking of Arboreal except for the Common Treeshrew *T*. *glis*, which is ranked as Scansorial. For our non-climbing species, we selected the Four-toed Hedgehog *Atelerix albiventris*, the Gray Short-tailed Opossum *Monodelphis domestica*, the Hispaniolan Solenodon *Solenodon paradoxus*, Trowbridge’s Shrew *Sorex trowbridgii*, and Botta’s Pocket Gopher *Thomomys bottae*. These taxa represent a variety of morphologies and a range of non-climbing locomotor modes, from ambulatory (Four-toed Hedgehog *A. albiventris*) to subterranean (Botta’s Pocket Gopher *T*. *bottae*). The Four-toed Hedgehog *A. albiventris* and Gray Short-tailed Opossum *M*. *domestica* are ranked as Terrestrial and the other three species are ranked at Inter-substrate.

### 3.6 Summary of Extant Mammal Results

Our binary and ordinal prediction results, presented in Figures 3 and 4, highlight three important properties of these models: the phylogenetic group-level effect strongly influences predictions, not every metric performs equally well within each taxon, and the multiple regression models greatly improve the predictive accuracy. We will address each of these properties individually.

**Figure 3:**
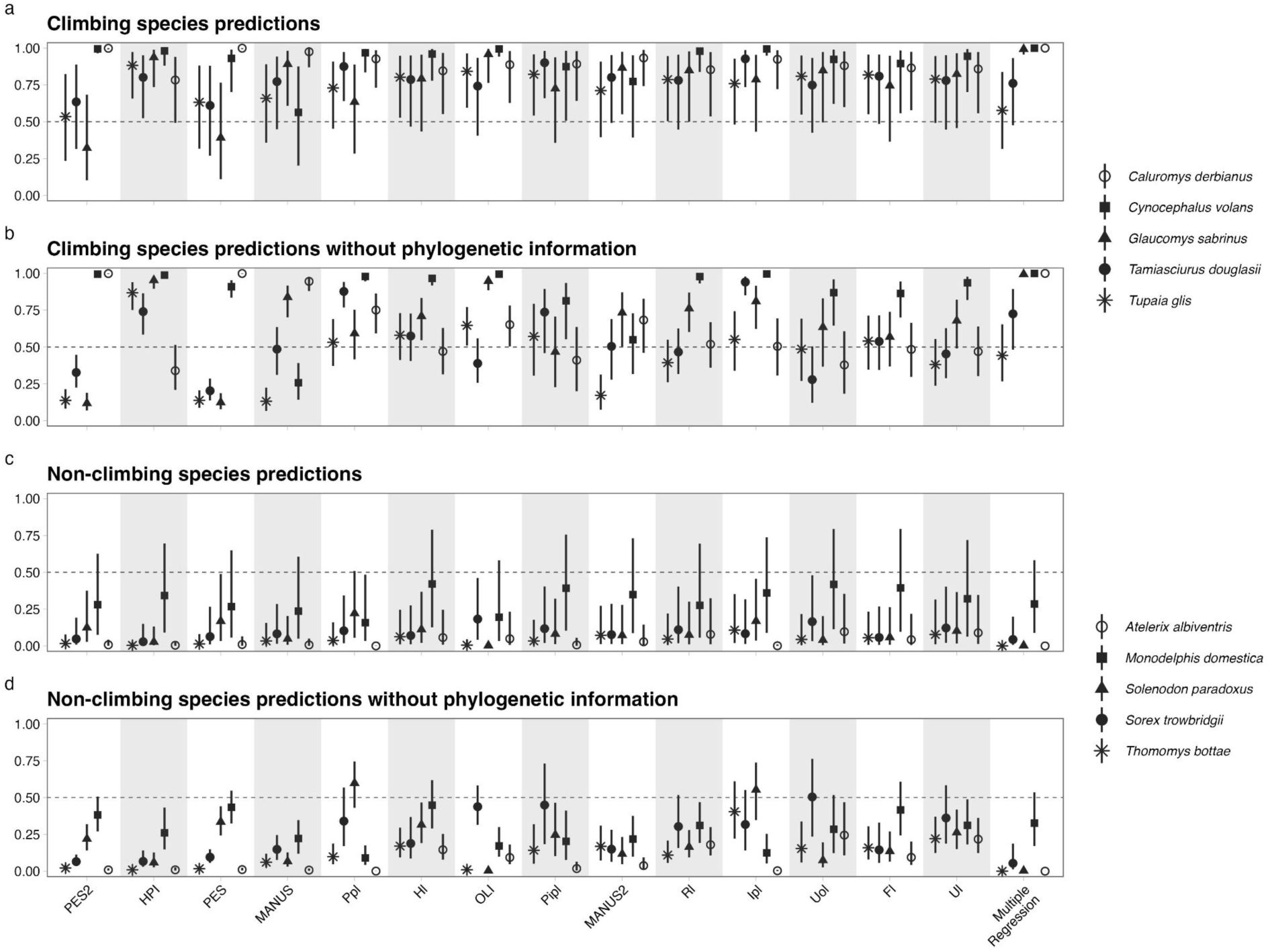
Predictions of binary climbing behavior in the “model” extant taxa both with and without the phylogenetic correlation matrix included. The 1 on the y-axis means 100% probability of climbing, and the 0 means 0% probability of climbing. The point-intervals depict the median and 89% probability distributions. Note that the climbing signal often moves toward the 50% probability line (dashed horizontal line, equal to a coin flip) when the phylogeny is not included. Also note that the multiple regression model predictions on the far right are quite similar with and without the phylogeny.

**Figure 4.**
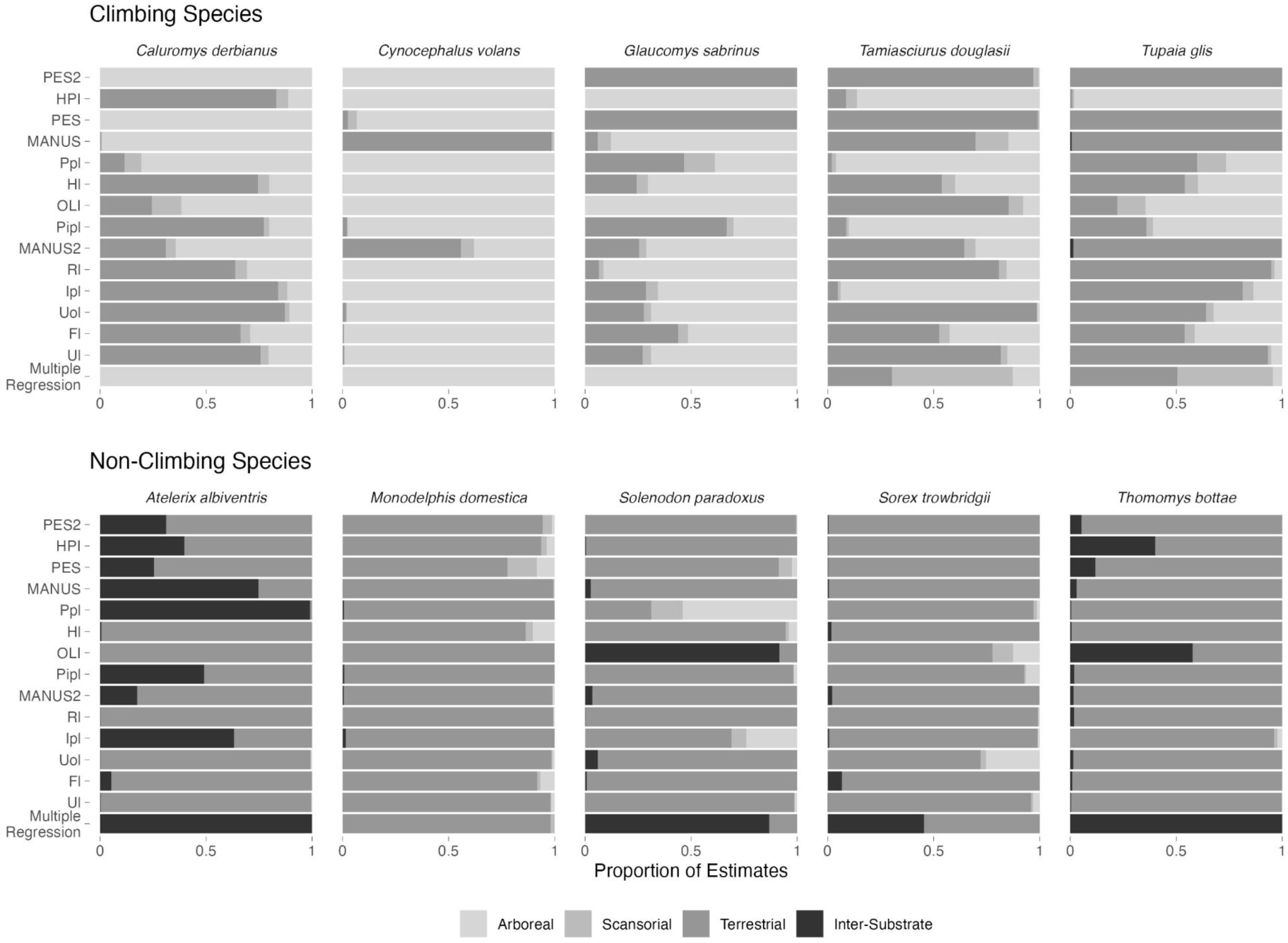
Predictions of ordinal locomotor rankings in the extant “model” taxa without the phylogenetic information. Bar plots depict the portion of the posterior draws that predict each locomotor rank. All climbing species have an ordinal ranking of Arboreal except the Common Treeshrew *T*. *glis*, which is ranked as scansorial. Among non-climbing species, The Four-toed Hedgehog *A. albiventris* and Gray Short-tailed Opossum *M*. *domestica* are ranked as terrestrial and the other three species are ranked at inter-substrate.

#### 3.6.1 The Effect of Phylogenetic Correlations on the Predictions

The extant species climbing predictions all demonstrate a very high level of accuracy (94–97%) when using the phylogenetic correlation matrix in the prediction. These results can be seen in Figure 3. When the predictions are made without the phylogenetic term, the accuracy drops precipitously (50–85%) except when using multiple regression (see below), demonstrating the degree of phylogenetic signal across the dataset. In other words, knowing both the morphological metric value and the phylogenetic position greatly improves prediction. These results suggest that predicting the locomotion of fossil taxa using individual traits in the absence of phylogenetic context may not be effective. Each clade of extant mammals has its own distinguishing features, which may be reflected in one or more appendicular skeleton measurements. This is true across phylogenetic scales: marsupials have distinct features not found in placentals, whereas muroid rodents may have postcranial synapomorphies not found in sciurids (see next paragraph). Additionally, the importance of phylogeny in the single metric model predictions highlights the need for appropriate comparative taxa when predicting locomotion in fossil or poorly studied species.

#### 3.6.2 Climbing is Predicted by Different Metrics for Different Taxa

For a single taxon, morphological metrics often generate very different probabilities of climbing. When using the full prediction method that includes the phylogenetic term (Figs. 3, 4), the ratio of the manual proximal phalanx length to the metacarpal length (MANUS) predicts a relatively low probability of climbing for the Philippine Colugo *C. volans*, yet the length of the intermediate phalanx (Ipl) and the ratio of pedal phalanges (PES, PES2) predict a nearly 100% probability of climbing. Similar but opposite, the two pedal phalanx ratios predict a toss-up between climbing and no climbing in the two climbing squirrel species (*G. sabrinus* and *T. douglasii*) and the Common Treeshrew *T. glis*, but most other metrics predict a high probability of climbing. The reasons for these discrepancies are due to some idiosyncratic morphologies of these species. Colugos are unusual for climbing mammals in that their intermediate manual phalanges are longer than their proximal phalanges (only one additional climbing species in our dataset has longer intermediate than proximal phalanges, the marsupial Brown Antechinus *Antichinus stuartii*), which may be related to roosting behavior (e.g., Meng et al., 2017). Similarly, for their body size, the Southern Flying Squirrel *G. sabrinus*, the Douglas’s Squirrel *T. douglasii*, and the Common Treeshrew *T. glis* have some of the longest metatarsals of any climbing species, which lowers their climbing predictions for the pedal phalanx ratios. Were we to make plots like Figure 3 for every species in the data set, we would likely find many more cases of specific metrics standing out as poor predictors for individual taxa, even among the most accurate measurements, demonstrating the risk of relying on a single measurement for prediction. Which brings us to the third observed property of our models.

#### 3.6.3 Multiple Regression Greatly Improves the Predictive Accuracy

When all the 14 best metrics are used together in a single model, our binary predictions show ∼95% accuracy with or without the phylogenetic covariance matrix included. For the non-phylogenetic predictions, this is a large improvement over 85% for PES2 (the best-performing individual metric) and provides an excellent modeling tool for fossil predictions. Our ordinal prediction results without phylogenetic information largely mirror the binary prediction results. As Figure 4 shows, locomotor ranking predictions vary among measurements, yet the multiple regression analysis that uses all 14 metrics performs quite well. Among the climbing taxa, all have a ranking of Arboreal except the Common Treeshrew *T. glis* which has a ranking of Scansorial. Using multiple regression, both *T. glis* and the Douglas’s Squirrel *Tamiasciurus douglasii* have a higher probability of being Scansorial than Arboreal, which reflects the reality that these two species do spend more time on the ground than the other three climbing taxa. Among the non-climbing taxa, most predictions match the true locomotor behavior of the species, with the Four-toed Hedgehog *Atelerix albiventris*, which we labeled Terrestrial, being the exception.

Digging deeper, the multiple regression results highlight the complexity of the sizes and proportions of appendicular skeletal elements as they relate to morphology. Looking at the binary and ordinal predictions for our model taxa (Figs. 3, 4), we can see that the multiple regression predictions cannot be attributed to a single metric or even several different metrics on their own. Put another way, the multiple regression predictions are not the same as averaging over the results from the 14 metrics. Rather, there is a complex interaction among the individual elements that the model can leverage but is difficult to interpret directly.

## 4 Linear Measurements as Robust Predictors of Arboreality

The goal of associating the size and shape of postcranial bones with a species’ ecology has a long and rich history. Linear measurements of limb elements, measures of size and proportion, have long dominated data-collection regimes. Measuring the length or width of a bone can be done with simple tools (rulers, calipers, photograph measurement tools) across a wide range of body sizes, and on bones in various states of preservation (loose bones, elements preserved in rock, X-ray images, etc.) in a relatively short amount of time. In recent decades, computational and imaging developments have led to a robust research field of shape analysis on 2D or 3D images, leading to advances in the fields of evolution and ecomorphology. 2D and 3D shape analyses are certainly powerful comparative tools appropriate for many questions, particularly when working with closely related taxa or species that occupy similar shape space (Zelditch et al., 2012), and they have successfully been used to differentiate mammalian locomotor modes (e.g., Janis and Martin-Serra, 2020). Nonetheless, our results demonstrate that linear measurements, functional indices, and log-shape ratios can (1) successfully identify features of the appendicular skeleton that are associated with climbing across the temporal and phylogenetic breadth of mammals, and (2) be used to estimate climbing can lead to highly accurate predictions. Thus, although more complex methodologies are useful for addressing certain questions, simple linear measurements are still robust and useful tools for inferring locomotor strategies, and they have the added benefits of being easily reproducible and posing a low barrier to entry.

### 4.1 Which Measurements Best Predict Climbing in Mammals?

Our results demonstrate that phenotypic measures associated with the hand and feet are the best predictors of climbing in mammals. Specifically, functional indices of the fingers and toes (PES, PES2, MANUS), log-transformed phalanx lengths, and LSRs of phalanx lengths all stand out as good predictors of climbing, a pattern noted in many previous studies (Lemelin, 1999; Kirk et al., 2008; Chen and Wilson, 2015; Nations et al., 2019; Weaver and Grossnickle, 2020). Intact hands and feet are quite rare in the fossil record, so it is worth noting that several limb measurements are effective predictors as well. In long bones, the humerus length, olecranon process length, ulna length, the humerus proximal width ratio (HPI), and the ratio of the olecranon process length to ulna length (OLI) offer the highest predictive power. Femur length and calcaneus length, either conditioned on body size or transformed as an LSR, both perform moderately well as predictors. A basic body plan of arboreal small mammals includes elongated forelimbs (closer to or equal in length to the hind limbs), gracile limb elements, and a shortened olecranon process of the ulna (e.g., Cartmill, 1985; Sargis, 2002a, 2002b; Salton and Sargis, 2008, 2009). Our long bone metrics support these findings.

#### 4.1.1 Pedal Metrics Predict Climbing as Well as or Better than Manual Metrics

Relative manual digit length has long been associated with locomotion in mammals, a functional relationship that our results confirm. Whereas many studies have focused on primates and primate origins (Bloch and Boyer, 2002; Gebo, 2004; Urbani and Youlatos, 2013), others have investigated relative finger length and locomotion across a range of mammalian taxa both living and extinct (Ji et al., 2002; Luo et al., 2003, Weisbecker and Warton, 2006; Luo, 2007; Sargis et al., 2007; Kirk et al., 2008; Luo et al., 2011; O’Leary et al., 2013; Meng et al., 2017, Nations et al 2019, Grossnickle et al 2020, Weaver and Grossnickle 2020). Though much attention has been focused on the hands, it is important to note that pedal digit length metrics perform equally well or better than finger lengths at predicting climbing (Fig. 2). In many mammal species, feet play an important functional role in grasping small-diameter substrates in horizontal movement (Cartmill, 1985), or in gripping substrates during vertical descent (Jenkins and McClearn, 1984), and may display adaptations for both (Youlatos et al., 2015; Meng et al., 2017). Our results suggest that, if available, pedal metrics should be included in any analysis of climbing.

#### 4.1.2 Single Metrics Can Effectively Predict Climbing… Some of the Time

Though most studies on the morphology of climbing in mammals focused on multiple skeletal measurements, individual measurements, indices, or ratios have on occasion been used to predict locomotion (e.g., Kirk et al., 2008). Exploring relationships between single metrics and locomotion is desirable, as it can provide a simple biological or functional morphological explanation (e.g., long fingers = climbing). Nations et al. (2019) found that the MANUS ratio effectively predicted locomotor rank in rodents using a three-tiered ranking system (Terrestrial, Climbing, Arboreal), and other studies found this index (or similar digit metrics) to be effective at predicting locomotion across mammals more generally (Kirk et al., 2008; Chen and Wilson, 2015; Weaver and Grossnickle, 2020).

Our results show that some single metrics do offer a way to estimate climbing in taxa with unknown locomotor affinity. As we noted above, manual and pedal phalanx ratios offer relatively high predictive accuracy as single metrics, even when predicting locomotion in species with unknown phylogenetic placement or lack of representation in the original model. However, our locomotor predictions of ‘model’ extant taxa demonstrate some potential pitfalls with using a single measurement (Fig 3). The primary problem is that clade-specific features can obscure true locomotor affinity when compared across a large phylogenetic breadth, such as the relatively long metatarsals found in squirrels, or the unusual manual phalanx ratios of colugos. Without adequate samples in the training dataset that match both the phenotype and the climbing rank, the single-metric predictions may be inaccurate.

An important caveat here is that our models were built using a large sample of phylogenetically diverse species from 70 families (42% of all mammalian families) representing most major mammal clades, including the monotreme “outgroup.” This is likely the reason that the predictions using the phylogenetic correlation matrix as a group-level effect were so accurate, despite no apparent overfitting, and the reason that prediction accuracy dropped substantially when predicting without the phylogeny (Figs. 1, 3). It is worth considering that models built for predictions at a narrower phylogenetic scale, such as predicting the locomotion of an extinct metatherian using data from living marsupials, may better account for clade-specific idiosyncrasies, thereby making single predictors more effective (a pattern observed in Nations et al. 2019).

#### 4.1.3 Using Multiple Predictors Improves Accuracy, but Limits Interpretation

Adding multiple metrics to our regression models greatly improves the accuracy of predictions. When using all 14 of our high-accuracy measurements to build the models, along with the intraspecific and phylogenetic group-level terms, the accuracy jumps to 95%, even without using the phylogenetic term in the prediction step. We view this as an acceptable range of accuracy for a model and are reasonably confident in our fossil predictions because of it. However, using more than one predictor variable in the models comes at a cost: interpretability.

Using multiple regression, the influence of a single predictor is contingent on all the other predictors used, often stated as “a change in one predictor if the other predictor(s) is held constant”, or as “conditioning” on another predictor (McElreath, 2020). Our original linear metric models are themselves multiple regression because they include both a linear measurement metric and a body size predictor (either log(mass) or geometric mean). Holding body size constant and only estimating the influence of the linear measurement, such as ulna length, makes intuitive sense, as ulna length cannot be interpreted in the context of locomotion without knowing something of the animal’s size. And, from such a model, we find that climbing is associated with a longer ulna at a given body size, directly addressing a simple, functional prediction. However, parsing out the effect of ulna length on locomotion while conditioning on body size, pedal phalanx ratios, humerus length, and relative olecranon process length is not straightforward, and we do not attempt to interpret our multiple regression outputs in this way. Therein lies the tradeoff of interpretability and predictive accuracy in this type of analysis.

### 4.2 Arboreal Species Rarely Exhibit Arboreal Adaptations for All Limb Traits

Adding to the dilemma between using a single metric to predict locomotion or using multiple metrics is that a species’ metrics very rarely all predict the same locomotor mode (e.g., Fig. 4). We’ve referenced the peculiarities of squirrel feet and colugo digits above, but this finding appears to be relevant across most or all mammal species. This observed pattern—that arboreal taxa rarely exhibit arboreal adaptations for all limb traits—may reflect many-to-one-mapping of form to function (Lauder, 1995; Wainwright et al., 2005). That is, there are likely many varying functional morphologies that permit effective climbing ability in mammals. For example, squirrel species largely depend on claws to grip substrates, whereas primates, which have nails instead of claws, use digits to grip the substrate in the arboreal milieu, which explains differences in digit morphology between these clades (Cartmill, 1985). Similarly, Runestad and Ruff (1995) and Grossnickle et al. (2020) observed that although most gliding mammals had relatively long limbs (likely an adaptation to help maximize patagial surface area), glider clades evolved different limb adaptations. For instance, some glider clades have relatively long zeugopods and others have relatively long stylopods, with differences between clades likely related to where patagia attach to limbs (e.g., at the wrists vs. elbows). An important implication of this observation is that if limb traits are used to infer arboreality (or other locomotor modes) in taxa with unknown locomotor behaviors (e.g., fossils), then researchers should not expect to see consistent support for arboreality across all traits even in taxa that were indeed arboreal.

### 4.3 Bayesian Multilevel Modeling as a Tool to Predict Locomotion

Throughout the past century, researchers have used numerous qualitative and quantitative methods to link morphology with locomotion. Common methods include regressing a metric on a body size variable then documenting the location of the residuals by locomotor mode, plotting two metrics together to observe how locomotor categories are distributed in morphospace, and using analysis of variance in a regression framework to compare the metrics among locomotor modes. In recent decades, multivariate ordination approaches have become standard, including but not limited to discriminant function analysis (DFA), principal components analysis (PCA), and canonical variance analysis (CVA) (e.g., see Dunn, 2018, for a review). Loadings may then be described qualitatively, or the distributions of different locomotor modes in morphospace are compared using secondary analyses of ordination scores. Predictions have been conducted using most of these approaches, but DFAs and CVAs are the most common way to predict locomotion in extinct taxa (e.g., Elissamburu and Vizcaíno, 2004; Samuels and Van Valkenburgh, 2008; Chen and Wilson, 2015; Meng et al., 2017; Grossnickle et al., 2020). Incorporating the evolutionary covariance of phenotypes generated by shared phylogenetic history can be included in many of these approaches, such as phylogenetic PCA, phylogenetic ANOVA/MANOVA, phylogenetic flexible discriminant analysis, and phylogenetic least squares regression (e.g., Schmitz and Motani, 2011; Polly et al., 2013; Adams and Collyer, 2018).

Here we take a less common approach to predicting climbing in extinct mammals that leverages the flexibility of multilevel modeling and Bayesian analysis. One key difference that plays out in multiple ways is the ability to estimate intraspecific variation along with phylogenetic covariance. The models in this study use the species term as a group-level effect, which estimates the effect size mean and variance within each species using partial pooling, and then uses these species-level estimates to generate the global prediction. This method integrates intraspecific variation into the analyses, and partial pooling removes any biases from uneven sampling among taxa (McElreath, 2020). In practice, this means that all the data collected can be used in the phylogenetic regression, regardless of the number of samples from each species (Hadfield and Nakagawa, 2010). This is non-trivial. Most modern phylogenetic comparative methods require that each tip in the phylogenetic tree is represented by a single data point, forcing users to take an average for each species. In our case, this would mean cutting our dataset nearly in half (427 samples down to 236; however, much more significant data loss is common in the literature), leading to fewer data points while simultaneously eliminating the biological reality of intraspecific variation from the analysis. Additionally, the effect of the phylogeny can directly be compared to the effect of intraspecific variation to determine which has a larger influence on the distribution of traits, a method common in quantitative genetics that is not often found in macroevolutionary analyses (de Villemereuil and Nakagawa, 2014).

An additional benefit of Bayesian regression is the ability to handle missing data in a probabilistic manner. Missing data are a reality of evolutionary biology, from skeletal elements missing in museum drawers to the inevitable incompleteness of fossil specimens. Our approach estimates missing values by leveraging the data that are complete (in our case, 12 linear measurements, body size, species membership, and the phylogenetic position) to estimate a posterior distribution of probable values for the missing points, while jointly incorporating these estimates into the larger climbing model. This method allows us to keep all the samples in the model training data to improve estimates.

Lastly, generalized linear modeling like logistic regression (binary response variables) and ordinal regression (rankings as response variables) depicts the morphology of climbing in two different but complementary contexts. Our binary models simply predict whether a species is known to climb, limiting the categorization to a single metric rather than using numerous descriptive categories of locomotion, and offering interpretable probabilities of climbing behavior between zero and one. Ordinal modeling of ranked locomotor preferences, on a scale of 1–4 (Inter-Substrate to Arboreal) incorporates the continuous nature of climbing behavior in mammals but does not treat the distances between ranks as fixed the way linear regression does for continuous points, nor does it treat each ranked preference as fully independent on the morphological axes, like a categorical analysis. The ordinal predictions do, however, allow us to interpret locomotor preference into a “categorical context” that tracks with a century of research using categorical descriptions of locomotion.

In sum, we believe that the methods presented here offer a well-grounded probabilistic framework to generate locomotor predictions in mammals. We should note that the methods presented are not limited to mammals or to locomotion. The models are flexible and can be easily modified to evaluate the importance of phenotypic characters to behavior, and to generate easy-to-interpret probabilities of any given ecological trait among taxa with poorly documented or unknown natural history.

## 5 Predicting Climbing in Fossil Mammals

We tested these new predictive models on extinct lineages from three clades of early mammals that have previously been interpreted to be mostly arboreal: Mesozoic Theria, Multituberculata, and Plesiadapiformes. In most instances, previous studies have used either comparative functional morphology, multivariate morphometrics, or some combination of the two, to infer the locomotor habits of these taxa. Except for some of the multituberculates, most of the taxa in our sample have previously been interpreted as being arboreal or scansorial. Thus, in addition to using them as case studies for demonstrating our approach to predicting arboreality, we can test previous locomotor hypotheses for these taxa.

### 5.1 Sampling of Fossil Mammals

We compiled skeletal measurements of extinct representatives of Multituberculata, Theria (we specifically sampled the earliest known therians with postcranial material), and Plesiadapiformes (i.e., stem primates). Measurements include values reported in the primary literature (available in Table S5) and were supplemented with measurements collected using photographs and scale bars (via *ImageJ*; Abràmoff et al., 2004). The fossil species include varying levels of missing data due to differences in the preservation of limb elements (Table S5).

As discussed in Section 2.2, multituberculates are often recovered in phylogenetic analyses as early crown mammals (stem therians), and they exhibit a diversity of locomotor modes (Simpson and Elftman, 1928; Jenkins and Krause, 1983; Luo et al., 2016; Kielan-Jaworowska and Qi, 1990; Kielan-Jaworowska and Gambaryan, 1994; Chen and Wilson, 2015; Weaver et al., 2021). The taxa included in our sample of multituberculates include taxa that have been interpreted as arboreal (*Ptilodus kummae*; Jenkins and Krause, 1983; Krause and Jenkins, 1983), scansorial–terrestrial (*Jeholbaatar kielanae*, *Rugosodon eurasiaticus*; Yuan et al., 2013; Chen and Wilson, 2015; Wang et al., 2019), terrestrial–saltatorial (*Kryptobaatar dashzevegi*; Kielan-Jaworowska and Gambaryan, 1994), and terrestrial–semifossorial (*Filikomys primaevus*; Weaver et al., 2021).

Our Mesozoic therians include one Jurassic species (*Juramaia sinensis*; Luo et al., 2011) and four Early Cretaceous species (*Eomaia scansoria*; Ji et al., 2002; *Sinodelphys szalayi*; Luo et al., 2003; *Amblolestes zhoui*, Bi et al., 2018; *Cokotherium jiufotangensis*, Wang et al., 2022), and they represent the earliest known therian skeletons. These taxa are most commonly classified as early eutherians, but we refer to them as therians because of their phylogenetic uncertainty. For instance, *Sinodelphys* was originally recovered in phylogenetic analyses as a metatherian (Luo et al., 2003; Huttenlocker et al., 2018), but additional morphological information published in Bi et al. (2018) led to *Sinodelphys* being recovered as a eutherian. Mao et al. (2020) incorporated the new morphological information from Bi et al. (2018) and still recovered *Sinodelphys* as a metatherian. Thus, there remains uncertainty about the phylogenetic position of *Sinodelphys*. Further, it is possible that some (or all) of these taxa are stem therians rather than crown therians (or eutherians)—they have been recovered as stem therians in some phylogenetic analyses (O’Leary et al., 2013; King and Beck, 2020, see their supplemental analyses; Krause et al., 2020). Nonetheless, we follow the most recent phylogenetic analyses in classifying the fossils as early therians (e.g., Huttenlocker et al., 2018; Krause et al., 2020; Mao et al., 2020; King and Beck, 2020). These early therians have been interpreted as either scansorial or arboreal (Ji et al., 2002; Luo et al., 2003; Luo et al., 2011; Chen and Wilson, 2015; Meng et al., 2017).

Our final fossil group are Plesiadapiformes, an early Paleogene group that first appear in the fossil record in the earliest Paleocene (Wilson Mantilla et al., 2021) and are likely stem primates (Silcox and Gunnell, 2008; Silcox et al., 2017). Our plesiadapiform sample includes representatives from four different families—Micromomyidae (*Dryomomys szalayi*), Palaechthonidae (*Torrejonia wilsoni*), Paromomyidae (*Ignacius clarkforkensis*), and Plesiadapidae (*Plesiadapis cookei*)—that span the middle–late Paleocene of North America. Plesiadapiforms have been almost universally interpreted as arboreal (Bloch et al. 2007; Bloch and Boyer 2007; Kirk et al. 2008; Chester et al. 2015; Chester et al. 2017), including the specimens analyzed here (Boyer and Gingerich, 2019; Chester et al., 2019).

### 5.2 Predictions of Fossil Mammal Locomotion

We generated binary and ordinal locomotor predictions for each of the 14 fossil taxa using the same methods as the extant taxa above, focusing specifically on the 14 metrics with the highest predictive accuracy (PES2, HPI, PES, MANUS, Ppl, Hl, OLI, Pipl, MANUS2, Rl, Ipl, Uol, Fl, and Ul), as well as multiple regression models. However, there are a few differences worth noting. A considerable number of our fossil taxa contain missing data due to incomplete preservation, and the missing measurements vary among the 14 taxa. Therefore, unlike the extant taxa, a single multiple regression model using all 14 covariates cannot generate predictions for every fossil taxon. To overcome this, we ran 10 additional multiple regression models that use the metrics available for each fossil taxon. For example, the multiple regression predictions for *Plesiadapis cookei* are from a model containing all 14 metrics, the *Sinodelphys* predictions are from a models that uses 11 metrics (PES, PES2, MANUS, MANUS2, Ppl, MANUS2, Ipl, Pipl, Hl, Rl, and Fl), and the predictions for *Torrejonia wilsoni* were generated from a model only containing two metrics (Ppl and Uol), the lowest number of any fossil taxon. Despite the limited number of measurements in some fossil taxa, the multiple regression model predictions still represent improvements in accuracy over the single predictor models as nine of the 11 multiple predictor models have an accuracy over 90% (Table S4).

#### 5.2.1 Results of Binary Fossil Predictions

Binary climbing predictions for fossil taxa varied among measurements and species, but most suggest reasonable estimates with acceptable amounts of uncertainty. All pedal ray ratio (PES or PES2) predictions, for the eight species that have those metrics available, suggest a very high probability of climbing; however, those predictions do not hold across most other metrics. For example, the early therians *Sinodephys* and *Eomaia* both have relatively long toes, as indicated by the high probability of climbing in the PES and PES2 metrics (Fig. 5), which suggests climbing, yet their radius and ulna lengths appear to be short, which indicates a low climbing probability in Hl and Rl (Fig 5). Interestingly, many of the early therians have similar predictions across all measurements, highlighting morphological features shared by these species. Relatively short radii, ulnae, and femora seem to be present in all the early therians, and the highest disparity is in the manual proximal phalanx metrics (MANUS, and Ppl).

**Figure 5:**
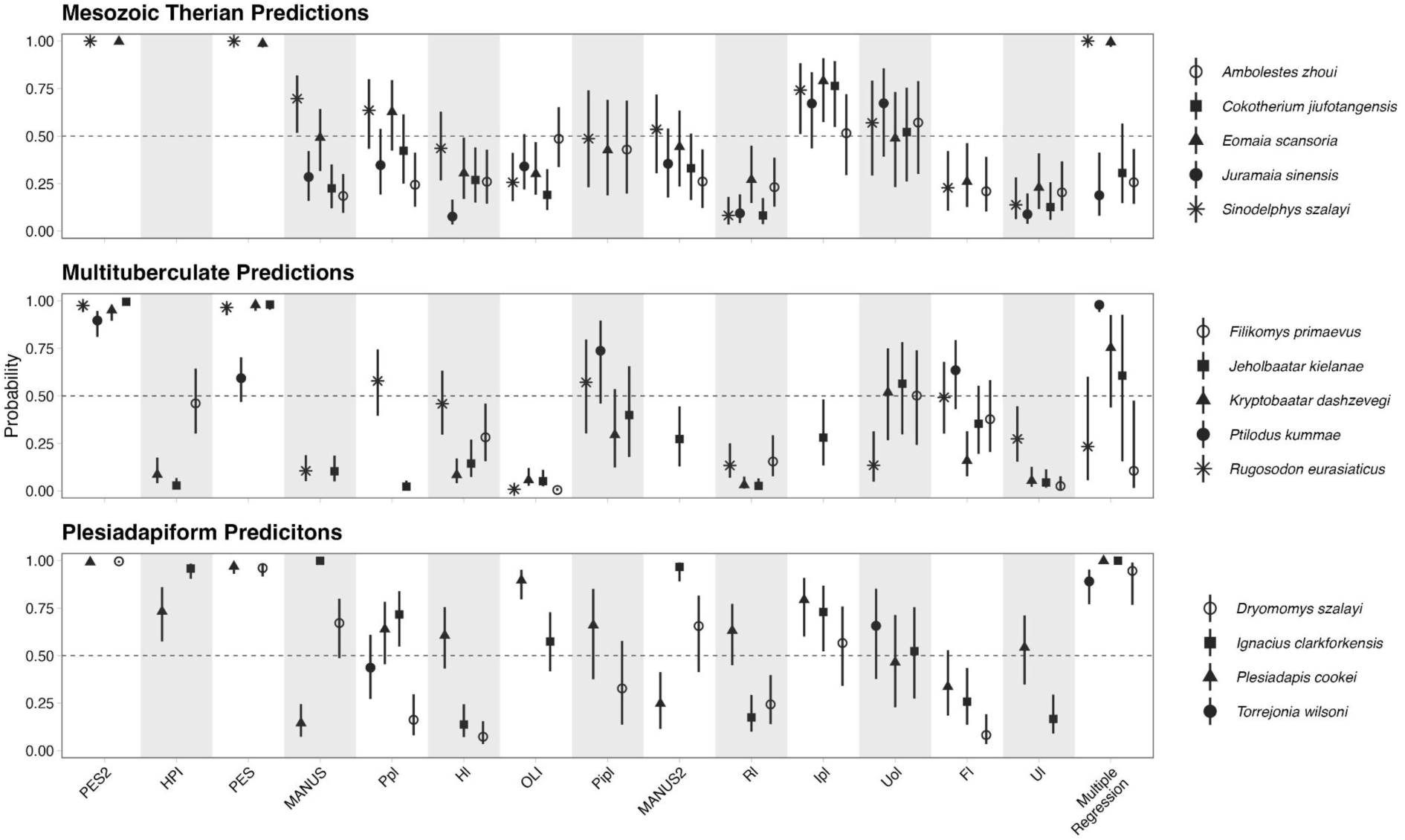
Predictions of binary climbing behavior in the fossil taxa. As with Figure 3, the 1 on the y-axis means 100% probability of climbing, and the 0 means 0% probability of climbing. The point intervals show the median prediction and the 89% probability distribution of each taxon for each metric.

The disparity in climbing predictions across metrics is greater among the multituberculates in our dataset than in the early therians, yet the disparity isn’t present for all metrics. Based on the length of most forelimb elements, the multituberculates are unlikely to do any climbing, although the length of the olecranon and the femur suggest more uncertainty. As with the early therian predictions, the pedal phalanges suggest a high probability of climbing for all multituberculates with those metrics available. Multiple regression predictions show substantial uncertainty in multituberculate climbing, and apart from *Ptilodus*, the predictions suggest anything from likely to climb to unlikely to climb, a coin-toss similar to our results for the Common Treeshrew *Tupaia glis* or the Gray Short-tailed Opossum *Monodelphis domestica*.

The plesiadapiform predictions are highly variable among metrics and species; however, the multiple regression prediction expresses confidence in climbing in all four taxa. This result highlights the complex structure of the multiple regression, where a specific suite of metric values at a specific body size can result in a prediction that does not match that of any individual element. Overall, the individual predictions have less structure across metrics in the plesiadapiforms than in early therians or multituberculates, such as a high variability in climbing predictions from manual ray metrics (MANUS, Ppl, etc.).

#### 5.2.2 Results of Ordinal Fossil Predictions

The ordinal predictions of the locomotor rank of the fossil taxa complement the binary predictions and offer a more concise way to discuss the locomotor habits of these species (Fig. 6). Three of the five Mesozoic therians, *Ambolestes*, *Cokotherium*, and *Juramaia* all have a robust probability of being ranked as Terrestrial based on the multiple regression results (Fig. 6). *Eomaia* is ranked as Scansorial or Arboreal, a similar prediction as the Douglas’s Squirrel and the Common Treeshrew. *Sinodelphys* has a high probability of being Arboreal. Like the binary predictions, the predicted ranks are not uniform across all metrics (although *Ambolestes* is close). *Eomaia* and *Sinodelphys* have Arboreal predictions for pedal phalanx ratios, measurements that the other early therians do not have. Although this may seem to affect the multiple regression predictions, they also are more scansorial or arboreal in most of the other 13 metric predictions than the other three taxa, suggesting that they would be ranked as Scansorial or Arboreal without the pedal ratios. The moderate probability of climbing predicted for all taxa in the olecranon process length does not have much influence on the multiple regression predictions.

**Figure 6.**
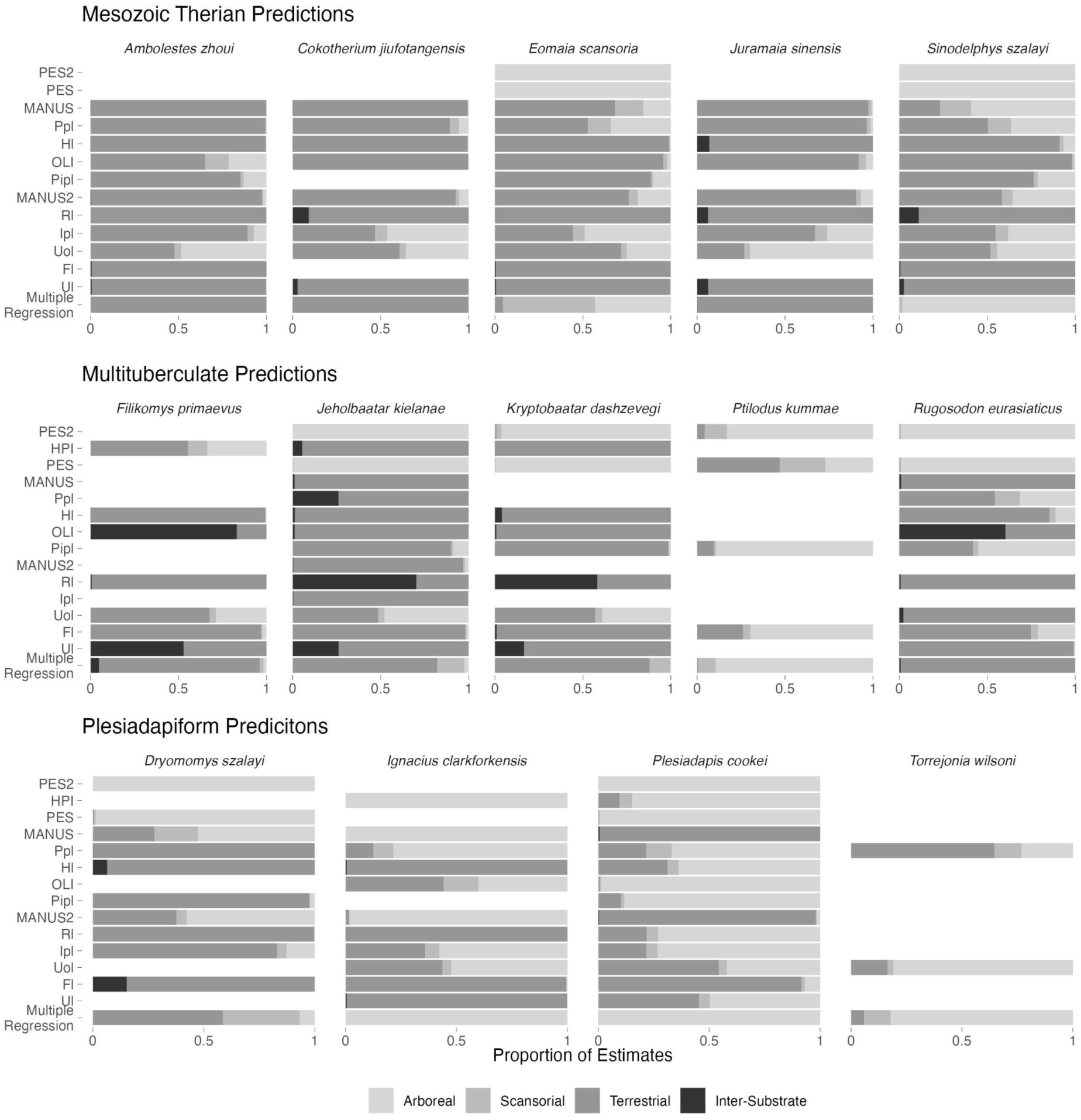
Predictions of ordinal locomotor rankings in the fossil taxa. Bar plots depict the portion of the posterior draws that predict each locomotor rank. Missing bars indicate missing metrics for each taxon.

The majority of multituberculate ordinal locomotion predictions are Terrestrial, but there is a small probability of scansoriality in *Jeholbaatar* and *Kryptobaatar*. Despite few available measurements from the 14 high-accuracy metrics, *Ptilodus* is strongly predicted to be arboreal, a pattern found across three of the four measurements and in the multiple regression prediction and consistent with previous studies (Jenkins and Krause, 1983; Krause and Jenkins, 1983). Unlike the Mesozoic therians, several multituberculate metrics suggest inter-substrate use, such as the relative olecranon length index (presumably quite large), and the radius length (presumably quite short).

The plesiadapiforms predictions appear to be mostly Terrestrial or Arboreal across all metrics, with the multiple regression providing a high probability of arboreality for all but *Dryomomys*, which has a split probability of Terrestrial and Scansorial. Many of the metrics provide very different predictions across species, such as MANUS2 ranging from 100% Arboreal to 100% Terrestrial, to an intermediate state in *Dryomomys* (Fig. 6).

## 6 Reevaluating the Importance of Arboreality in Mammalian Evolution

Here we have briefly reviewed the evolution of arboreality among Mesozoic and early Cenozoic mammals and, using a new approach compared to previous fossil studies, have re-examined the potential locomotor habits of three groups that have, either currently or historically, been considered to be mostly arboreal: Mesozoic Theria, Multituberculata, and Plesiadapiformes. This exercise has revealed an observation that, to our knowledge, has not been adequately highlighted in previous literature: arboreality may have been relatively uncommon among early mammals. Below, we explore the implications of this finding, derived from both our review of early mammalian arboreality in Section 2 and our new locomotor predictions in Section 5.

### 6.1 Were Therians Ancestrally Ground-Dwellers?

That Mammalia were ancestrally arboreal is a long-standing hypothesis in mammalian evolution (Matthew, 1904), yet most of the debates about ancestral mammalian locomotor strategies have centered on Metatheria and Eutheria (see Section 2) and occurred in a time when little was known about the postcranial morphologies of Mesozoic mammals. In the past 25 years, our knowledge of early mammalian locomotion has grown immensely, revealing a rich array of locomotor specializations that spanned swimmers and burrowers to gliders (Luo, 2007; Grossnickle et al., 2019). Despite the overall increase in known locomotor diversity, all of the recently discovered therian skeletons from the Jurassic and Early Cretaceous have been inferred to have had scansorial or arboreal locomotor modes (Ji et al., 2002; Luo et al., 2003; Luo et al., 2011; Chen and Wilson, 2015; Bi et al., 2018), reinforcing the hypothesis that arboreal locomotion was the ancestral condition for Theria (Matthew, 1904). Further, because arboreality is still proposed as the predominant locomotor mode among early therians, it has played an outsized role in our understanding of early mammalian evolution.

Our fossil predictions allow us to evaluate the hypothesis that therians are ancestrally arboreal. We rely on the multiple regression results for our primary fossil predictions because these results performed especially well, and they incorporate all of the best-performing predictors of climbing.

Our results conflict with some previous inferences of scansoriality–arboreality in early therians. Of the early therians included in our fossil sample, only *Eomaia* and *Sinodelphys* exhibited strong evidence of arboreality, whereas the remaining therians (*Ambolestes*, *Cokotherium*, and *Juramaia*) had a low probability of being climbers (most apparent in the ordinal results; Fig 6). Most notable of those taxa is *Juramaia*, which is the temporally oldest (by ca. 30 million years) and likely earliest branching of our sampled therians, meaning that it is most likely among our sample to represent the ancestral state of therians. Luo et al. (2011) inferred that *Juramaia* was scansorial based on proportional lengths of manual bones, via comparisons to those of extant mammals (Kirk et al., 2008). As we emphasized in Section 5.2, locomotor estimates derived from a single portion of the appendicular skeleton can be misleading. By incorporating data from across the appendicular skeleton, our regression models predict that *Juramaia* was terrestrial. This suggests that therians could be ancestrally terrestrial, not scansorial or arboreal.

Locomotor inferences for stem therians provide additional evidence that therians (and possibly mammals in general) were ancestrally terrestrial. As we review in Section 2, most of synapsid evolution is characterized by terrestrial and semifossorial locomotion, so it is likely that ground-dwelling is the ancestral mammalian locomotor strategy. This is evident in the postcranial morphology of both non-mammalian mammaliaforms and early crown mammals, many of which exhibit forelimb structures that belie a propensity for scratch-digging (e.g., Luo and Wible, 2005; Luo et al., 2015; Panciroli et al., 2021; Weaver et al., 2021; but note that one cladotherian, *Henkelotherium*, has been inferred to be scansorial [Jäger et al., 2020]). Although morganucodontans are less extreme in their postcranial adaptations for digging (e.g., Jenkins and Parrington, 1976), they too lack evidence of specialized climbing abilities and were likely, at least, terrestrial (Chen and Wilson, 2015). Taken together, the evidence suggests that ground-dwelling locomotion was common among the phylogenetic backbone of early mammals, with arboreality being a derived behavior that evolved in many early mammal groups independently.

An important caveat here is that the early therians in our sample are often classified as eutherians (e.g., Luo et al., 2011, Bi et al., 2018). If this is the case, our results could suggest that *eutherians*, but not necessarily *therians*, were ancestrally terrestrial. Such an interpretation is in line with previous studies that posited that eutherians were ancestrally terrestrial and metatherians were ancestrally arboreal (Huxley, 1880; Bensley, 1903; Dollo, 1899; Haines, 1958; Szalay, 1981; Szalay, 1984; Argot, 2001; Argot, 2002; Williamson et al., 2014). Unfortunately, we cannot easily test this hypothesis about the eutherian–metatherian dichotomy in locomotor mode because early metatherian skeletons are lacking in the fossil record. Indeed, the oldest unequivocal metatherian fossils are from the middle Cretaceous of North America (Davis et al., 2008; Bi et al., 2018) and are only represented by teeth. The only associated, relatively complete, and unequivocal metatherian skeleton comes from the Late Cretaceous (Campanian) of Mongolia (*Asiatherium*; Trofimov and Szalay, 1994), and it has been interpreted to be an obligate terrestrialist (Szalay and Trofimov, 1996). All other Mesozoic metatherian postcranial remains are fragmentary, which complicates locomotor inferences (e.g., Szalay, 1994; Horovitz, 2000; Szalay and Sargis, 2006; Chester et al., 2010; Chester et al., 2012; DeBey and Wilson, 2014; DeBey and Wilson, 2017; Freimuth et al., 2021). Thus, it is possible that the earliest metatherians were indeed arboreal (*sensu* Szalay, 1981), but our earliest glimpse into their locomotor habits, like the putative eutherians discussed above, suggests a much more terrestrial locomotor mode. Ultimately, testing these competing hypotheses will require the discovery of more complete early metatherian specimens.

### 6.2 Many Mesozoic Paleoenvironments Promoted Terrestriality, Not Arboreality

Here, we highlighted that arboreal locomotion among early mammals, although likely evolving independently in multiple groups (see Section 2), may have been relatively rare compared to more terrestrial or semifossorial locomotor strategies. Indeed, in addition to the discussion of therian mammals above, only the youngest, Paleocene representative of multituberculates in our sample (*Ptilodus*) is inferred to have been truly arboreal (Figs 5 and 6). This prevalence of ground-dwelling is also consistent among other Mesozoic mammaliaform groups not included in our analyses (e.g., eutriconodontans, docodontans; see Section 2). Thus, we propose that (1) most Mesozoic paleoenvironments likely promoted terrestriality and semifossoriality among mammals, and (2) arboreality did not become prevalent among mammalian communities until the early Paleogene, concomitant with the rise of dense, angiosperm-dominated, closed-canopy forests.

Although often envisaged as lush tropical rainforests, most Mesozoic terrestrial ecosystems were likely characterized by a relatively open vegetation structure (e.g., Wing and Tiffney, 1987; Wing and Sues, 1992; Carvalho et al., 2021) and relatively strong seasonality (especially in precipitation; e.g., Fricke et al., 2010; Burgener et al., 2019). In extant terrestrial ecosystems, the ecomorphological structure of mammalian communities closely tracks these vegetational and climatic variables, with open and seasonally variable habitats being characterized by most terrestrial and burrowing taxa, rather than diverse tree-dwelling groups that characterize tropical, closed-canopy forests (e.g., Chen et al., 2019). Thus, it should be expected that Mesozoic mammals would be mostly ground-dwelling rather than tree-dwelling (e.g., Haines, 1958).

Consistent with the hypothesized prevalence of ground-dwelling adaptations among Mesozoic mammals is the exceptional Yanliao Biota of China, which mostly come from the Middle Jurassic Tiaojishan Formation (Xu et al., 2016; Li et al., 2023). Unlike the more open– patchy vegetation that likely characterized most Mesozoic paleoenvironments, the Tiaojishan Formation flora is indicative of a dense, temperate-to-subtropical forest hosting conifer trees with an average height of 25 m (Jiang et al., 2019). Accordingly, most of the arboreal mammals and mammaliaforms reported from the Mesozoic come from the Tiaojishan Formation, including the only known arboreal docodont (*Agilodocodon*; Meng et al., 2015) and one of the only known arboreal or gliding eutriconodontans (*Volaticotherium*; Meng et al., 2006). All of gliding ‘haramiyidans’, which as discussed above (Section 2.1) were the major exception to the scarcity of arboreal adaptations among non-therian mammaliaforms, are also from the Tiaojishan Formation, and a recent study suggested that tall-tree forests promote the evolution of gliding among vertebrates (Wagner et al., 2023). Whether this exception is specific to the Tiaojishan Formation or indicative of the Middle Jurassic more broadly is unclear, but there is evidence for more densely vegetated paleoenvironments elsewhere in the Middle Jurassic (e.g., Italy; Costamanga et al., 2017). In either case, where we have a clear exception to the ‘open-Mesozoic-habitats rule’ in the Tiaojishan Formation, we see a mammaliaform community that exhibits a much greater prevalence of arboreality, consistent with the hypothesis that pervasive arboreality is contingent on vegetation structure (at least in part).

Following the K-Pg mass extinction, the continued diversification of angiosperms led to the establishment of dense, closed-canopy vegetation that was comparable to modern tropical rainforests (e.g., Wing and Tiffney, 1987; Johnson and Ellis, 2002; Benton et al., 2021; Carvalho et al., 2021). Again, given expectations from modern mammalian communities, we would expect this rise of tropical forests to be accompanied by a rise in mammalian arboreality, and this is borne out in the fossil record. By the late Paleocene to early Eocene, arboreality may have been the most common mammalian locomotor strategy (e.g., Carvalho et al., 2021; Benton et al., 2022). Thus, although some Mesozoic mammals were certainly arboreal, most of those arboreal taxa were likely living in densely forested paleoenvironments that were functionally analogous to temperate–subtropical rainforests today. In contrast, most Mesozoic mammals were likely living in much more open or patchily vegetated (e.g., Wing and Tiffney, 1987; Wing and Sues, 1992; Carvalho et al., 2021) and seasonal (Fricke et al., 2010; Burgener et al., 2019) paleoenvironments. As such, we propose that terrestriality and semifossoriality were the most common early mammalian locomotor strategies, and the shift to predominant arboreality did not occur until after the K-Pg boundary.

## Conclusion

We provided an overview of mammal arboreality through the fossil record, and presented some novel methods that we believe offer a solid probabilistic framework for any researcher to predict locomotion in synapsids from the Paleozoic to the present. Based on our review of arboreality among early mammals and our analysis of arboreality among representative early mammalian groups, we suggest that arboreality, although having evolved independently in many early mammal lineages, was not as important to the early evolutionary history of mammals as has been previously proposed. Instead, we suggest that most early mammals, including the ancestors to crown Theria, were ground-dwellers, which is consistent with our understanding of Mesozoic paleoenvironments as much more seasonal and characterized by open or patchy vegetation structure. To the authors, the conclusion that the ancestor of modern mammals was not arboreal makes the repeated evolution of arboreality and the correlated phenotypic adaptations across so many mammal clades even more fascinating. That arboreality has featured so prominently in the story of mammalian evolution is likely due, at least in part, to the prominence of arboreality in the evolutionary history of our own lineages, Primates. The major features that make primates primates—big brains, stereoscopic vision, opposable thumbs, and a propensity for sweets—have all been attributed to a life in the trees. However, our ancient mammalian roots were likely planted in the ground, which provides a new lens through which we can view the evolution of key mammalian adaptations. Just as we have now come to view primate evolution from the vantage of a tree, perhaps we can now begin to peer into the lives of the earliest mammals through the opening of a burrow.

## ACKNOWLEDGMENTS

For feedback and assistance with data collection, we thank Chris Law, Kathryn Stanchak, Stephanie Smith, Spencer Pevsner, Meng Chen, Gregory Wilson Mantilla, Richard Meyn, Kenneth-Dieter Benton. We also thank two anonymous reviewers for helpful suggestions that greatly improved our study.

## Footnotes

1 See discussion at https://discourse.mc-stan.org/t/predict-with-a-brms-phylogenetic-model-for-a-new-species-with-known-phylogenetic-position/7480/8

## LITERATURE CITED

Abràmoff, M. D., Magalhães, P. J., & Ram, S. J. (2004). Image processing with ImageJ. Biophotonics International, 11(7), 36–42.

Adams, D. C., & Collyer, M. L. (2018). Phylogenetic ANOVA: Group-Clade Aggregation, Biological Challenges, and a Refined Permutation Procedure. Evolution, 72(6), 1204–15.

Archibald, J. D. (1998). Archaic ungulates (“Condylarthra”). In: Janis CM, Scott KM, Jacobs LL (Eds.), Evolution of Tertiary mammals of North America. Vol. 1: Terrestrial carnivores, ungulates, and ungulatelike mammals.

Argot, C. (2001). Functional-adaptive anatomy of the forelimb in the Didelphidae, and the paleobiology of the Paleocene marsupials Mayulestes ferox and Pucadelphys andinus. Journal of Morphology, 247(1), 51–79.

Argot, C. (2002). Functional-adaptive analysis of the hindlimb anatomy of extant marsupials and the paleobiology of the Paleocene marsupials Mayulestes ferox and Pucadelphys andinus. Journal of Morphology, 253(1), 76–108.

Argot, C. (2013). Postcranial analysis of a carnivoran-like archaic ungulate: the case of Arctocyon primaevus (Arctocyonidae, Mammalia) from the late Paleocene of France. Journal of Mammalian Evolution, 20, 83–114.

August, P. V. (1983). The role of habitat complexity and heterogeneity in structuring tropical mammal communities. Ecology, 64(6), 1495–1507.

Bensley, B. A. (1903). On the Evolution of the Australian Marsupialia; with Remarks on the Relationships of the Marsupials in general. Transactions of the Linnean Society of London. 2nd Series: Zoology, 9(3), 83–217.

Benton, M. J., Wilf, P., & Sauquet, H. (2022). The Angiosperm Terrestrial Revolution and the origins of modern biodiversity. New Phytologist, 233(5), 2017–2035.

Bi, S., Wang, Y., Guan, J., Sheng, X., & Meng, J. (2014). Three new Jurassic euharamiyidan species reinforce early divergence of mammals. Nature, 514(7524), 579–584.

Bi, S., Zheng, X., Wang, X., Cignetti, N. E., Yang, S., & Wible, J. R. (2018). An Early Cretaceous eutherian and the placental–marsupial dichotomy. Nature, 558(7710), 390–395.

Bloch, J. I., & Boyer, D. M. (2002). Grasping Primate Origins. Science, 298(5598), 1606–1610.

Bloch, J. I., Silcox, M. T., Boyer, D. M., & Sargis, E. J. (2007). New Paleocene skeletons and the relationship of plesiadapiforms to crown-clade primates. Proceedings of the National Academy of Sciences, 104(4), 1159–1164.

Bloch, J. I., & Boyer, D. M. (2007). New skeletons of Paleocene-Eocene Plesiadapiformes: a diversity of arboreal positional behaviors in early primates. In Primate origins: adaptations and evolution (pp. 535–581). Boston, MA: Springer US.

Boyer, D. M., & Gingerich, P. D. (2019). Skeleton of late Paleocene *Plesiadapis cookei* (Mammalia, Euarchonta): Life history, locomotion, and phylogenetic relationships.

Burgener, L., Hyland, E., Huntington, K. W., Kelson, J. R., & Sewall, J. O. (2019). Revisiting the equable climate problem during the Late Cretaceous greenhouse using paleosol carbonate clumped isotope temperatures from the Campanian of the Western Interior Basin, USA. Palaeogeography, Palaeoclimatology, Palaeoecology, 516, 244–267.

Bürkner, P. C. (2018). Advanced Bayesian Multilevel Modeling with the R Package Brms. The R Journal, 10(1), 395–411. doi: 10.32614/RJ-2018-017

Burtner, A. E., Grossnickle, D. M., Santana, S. E., & Law, C. J. (2024). Gliding toward an understanding of the origin of flight in bats. PeerJ, 12, e17824.

Calede, J. J., Samuels, J. X., & Chen, M. (2019). Locomotory adaptations in entoptychine gophers (Rodentia: Geomyidae) and the mosaic evolution of fossoriality. Journal of Morphology, 280(6), 879–907.

Cartmill, M. (1985). Climbing. In Functional Vertebrate Morphology, edited by M. Hildebrand, D. M. Bramble, K. F. Liem, and Wake D. B, 73–88. Cambridge: Harvard University Press.

Carvalho, M. R., Jaramillo, C., de la Parra, F., Caballero-Rodríguez, D., Herrera, F., Wing, S., … Martinez, C. (2021). Extinction at the end-Cretaceous and the origin of modern Neotropical rainforests. Science, 372(6537), 63–68.

Chen, M., & Luo, Z. X. (2013). Postcranial skeleton of the Cretaceous mammal *Akidolestes cifellii* and its locomotor adaptations. Journal of Mammalian Evolution, 20, 159–189.

Chen, M., & Wilson, G. P. (2015). A multivariate approach to infer locomotor modes in Mesozoic mammals. Paleobiology, 41(2), 280–312.

Chen, M., Strömberg, C. A., & Wilson, G. P. (2019). Assembly of modern mammal community structure driven by Late Cretaceous dental evolution, rise of flowering plants, and dinosaur demise. Proceedings of the National Academy of Sciences, 116(20), 9931–9940.

Chester, S. G., Sargis, E. J., Szalay, F. S., Archibald, J. D., & Averianov, A. O. (2010). Mammalian distal humeri from the Late Cretaceous of Uzbekistan. Acta Palaeontologica Polonica, 55(2), 199–211.

Chester, S. G., Sargis, E. J., Szalay, F. S., Archibald, J. D., & Averianov, A. O. (2012). Therian femora from the Late Cretaceous of Uzbekistan. Acta Palaeontologica Polonica, 57(1), 53–64.

Chester, S. G., Bloch, J. I., Boyer, D. M., & Clemens, W. A. (2015). Oldest known euarchontan tarsals and affinities of Paleocene *Purgatorius* to Primates. Proceedings of the National Academy of Sciences, 112(5), 1487–1492.

Chester, S. G., Williamson, T. E., Bloch, J. I., Silcox, M. T., & Sargis, E. J. (2017). Oldest skeleton of a plesiadapiform provides additional evidence for an exclusively arboreal radiation of stem primates in the Palaeocene. Royal Society Open Science, 4(5), 170329.

Chester, S. G., Williamson, T. E., Silcox, M. T., Bloch, J. I., & Sargis, E. J. (2019). Skeletal morphology of the early Paleocene plesiadapiform *Torrejonia wilsoni* (Euarchonta, Palaechthonidae). Journal of Human Evolution, 128, 76–92.

Cinelli, C., Forney, A., & Pearl, J. (2022). A Crash Course in Good and Bad Controls. Sociological Methods & Research. 0 10.1177/00491241221099552

Claude, J. (2013). Log-shape ratios, Procrustes superimposition, elliptic Fourier analysis: three worked examples in R. Hystrix, 24, 94–102.

Collinson, M. E., & Hooker, J. J. (1991). Fossil evidence of interactions between plants and plant-eating mammals. Philosophical Transactions of the Royal Society of London. Series B: Biological Sciences, 333(1267), 197–208.

Costamagna, L. G., Kustatscher, E., Scanu, G. G., Del Rio, M., Pittau, P., & van Konijnenburg-van Cittert, J. H. (2018). A palaeoenvironmental reconstruction of the Middle Jurassic of Sardinia (Italy) based on integrated palaeobotanical, palynological and lithofacies data assessment. Palaeobiodiversity and Palaeoenvironments, 98, 111–138.

Davis, B. M., Cifelli, R. L., & Kielan-Jaworowska, Z. (2008). Earliest evidence of Deltatheroida (Mammalia: Metatheria) from the Early Cretaceous of North America. Mammalian evolutionary morphology: a tribute to Frederick S. Szalay, 3-24.

de Villemereuil, P., & Nakagawa, S. (2014). General quantitative genetic methods for comparative biology. In Modern phylogenetic comparative methods and their application in evolutionary biology: concepts and practice (pp. 287–303).

DeBey, L. B., & Wilson, G. P. (2014). Mammalian femora across the Cretaceous–Paleogene boundary in eastern Montana. Cretaceous Research, 51, 361–385.

DeBey, L. B., & Wilson, G. P. (2017). Mammalian distal humerus fossils from eastern Montana, USA with implications for the Cretaceous-Paleogene mass extinction and the adaptive radiation of placentals. Palaeontologia Electronica, 20(3), 1–93.

Deischl, D.G. (1964). The Postcranial Anatomy of Cretaceous Multituberculate Mammals. M.Sc. Thesis. University of Minnesota, Minneapolis, 85 pp.

Dollo, L. (1899). Les ancêtres des Marsupiaux étaient-ils arboricoles?. Trav. Stat. Zool., Wimereux, 7, 188–203.

Dunn, R. H. (2018). Functional morphology of the postcranial skeleton. In Methods in paleoecology: Reconstructing Cenozoic terrestrial environments and ecological communities (pp. 23–36).

Elissamburu, A., & Vizcaíno, S. F. (2004). Limb proportions and adaptations in caviomorph rodents (*Rodentia: Caviomorpha*). Journal of Zoology, 262(2), 145–159.

Eriksson, O. (2016). Evolution of angiosperm seed disperser mutualisms: the timing of origins and their consequences for coevolutionary interactions between angiosperms and frugivores. Biological Reviews, 91(1), 168–186.

Freckleton, R. P. (2009). The seven deadly sins of comparative analysis. Journal of Evolutionary Biology, 22(7), 1367–1375.

Freimuth, W. J., Varricchio, D. J., Brannick, A. L., Weaver, L. N., & Wilson Mantilla, G. P. (2021). Mammal-bearing gastric pellets potentially attributable to Troodon formosus at the Cretaceous Egg Mountain locality, Two Medicine Formation, Montana, USA. Palaeontology, 64(5), 699–725.

Fricke, H.C., Foreman, B.Z., & Sewall, J.O. (2010). Integrated climate model-oxygen isotope evidence for a North American monsoon during the Late Cretaceous. Earth and Planetary Science Letters, 289(1-2), 11–21.

Fröbisch, J., & Reisz, R. R. (2009). The Late Permian herbivore *Suminia* and the early evolution of arboreality in terrestrial vertebrate ecosystems. Proceedings of the Royal Society B: Biological Sciences, 276(1673), 3611–3618.

Fröbisch, J., & Reisz, R. R. (2011). The postcranial anatomy of *Suminia getmanovi* (Synapsida: Anomodontia), the earliest known arboreal tetrapod. Zoological Journal of the Linnean Society, 162(3), 661–698.

Gebo, D. L. (2004). A shrew-sized origin for primates. American Journal of Physical Anthropology, 47, 40–62.

Gelman, A., & Rubin, D. B. (1992). Inference from iterative simulation using multiple sequences. Statistical Science, 7(4), 457–511.

Gidley, J.W. (1909). Notes on the fossil mammalian genus Ptilodus, with descriptions of new species. Proceedings of the United States National Museum 36, 611–627.

Godinot, M., & Prasad, G.V.R. (1994). Discovery of Cretaceous arboreal eutherians. Naturwissenschaften, 81, 79–81.

Goswami, A., Prasad, G.V., Upchurch, P., Boyer, D.M., Seiffert, E.R., Verma, O., Gheerbrant, E., & Flynn, J.J. (2011). A radiation of arboreal basal eutherian mammals beginning in the Late Cretaceous of India. Proceedings of the National Academy of Sciences, 108(39), 16333–16338.

Gould, F. D., & Rose, K. D. (2014). Gnathic and postcranial skeleton of the largest known arctocyonid ‘condylarth’ *Arctocyon mumak* (Mammalia, Procreodi) and ecomorphological diversity in Procreodi. Journal of Vertebrate Paleontology, 34(5), 1180–1202.

Granatosky, M. C. (2018). A review of locomotor diversity in mammals with analyses exploring the influence of substrate use, body mass and intermembral index in primates. Journal of Zoology, 306(4), 207–216.

Grossnickle, D. M., Smith, S. M., & Wilson, G. P. (2019). Untangling the multiple ecological radiations of early mammals. Trends in Ecology & Evolution, 34(10), 936–949.

Grossnickle, D. M., Chen, M., Wauer, J. G., Pevsner, S. K., Weaver, L. N., Meng, Q. J., …, & Luo, Z. X. (2020). Incomplete convergence of gliding mammal skeletons. Evolution, 74(12), 2662–2680.

Guignard, M. L., Martinelli, A. G., & Soares, M. B. (2019). The postcranial anatomy of *Brasilodon quadrangularis* and the acquisition of mammaliaform traits among non-mammaliaform cynodonts. PloS One, 14(5), e0216672.

Hadfield, J. D., & Nakagawa, S. (2010). General Quantitative Genetic Methods for Comparative Biology: Phylogenies, Taxonomies and Multi-Trait Models for Continuous and Categorical Characters. Journal of Evolutionary Biology, 23(3), 494–508.

Haines, R. W. (1958). Arboreal or terrestrial ancestry of placental mammals. The Quarterly Review of Biology, 33(1), 1–23.

Han, G., Mao, F., Bi, S., Wang, Y. and Meng, J. (2017). A Jurassic gliding euharamiyidan mammal with an ear of five auditory bones. Nature, 551(7681), 451–456.

Hauser, D. C. (1964). Anting by gray squirrels. Journal of Mammalogy, 45(1), 136–138.

Hedrick, B. P., Dickson, B. V., Dumont, E. R., & Pierce, S. E. (2020). The evolutionary diversity of locomotor innovation in rodents is not linked to proximal limb morphology. Scientific Reports, 10(1), 717.

Hellert, S. M., Grossnickle, D. M., Lloyd, G. T., Kammerer, C. F., & Angielczyk, K. D. (2023). Derived faunivores are the forerunners of major synapsid radiations. Nature Ecology & Evolution, 1-11.

Hildebrand, M. (1985A). Digging of quadrupeds. In: Hildebrand M, Bramble DM, Liem KF, Wake DB (Eds.), Functional Vertebrate Morphology. Cambridge: Harvard University Press. 89–109.

Hildebrand, M. (1985B). Walking and running. In: Hildebrand M, Bramble DM, Liem KF, Wake DB (Eds.), Functional Vertebrate Morphology. Cambridge: Harvard University Press. 38–57.

Hoffmann, S., Beck, R. M., Wible, J. R., Rougier, G. W., & Krause, D. W. (2020A). Phylogenetic placement of *Adalatherium hui* (Mammalia, Gondwanatheria) from the Late Cretaceous of Madagascar: implications for allotherian relationships. Journal of Vertebrate Paleontology, 40(sup1), 213–234.

Hoffmann, S., Hu, Y., & Krause, D. W. (2020B). Postcranial morphology of *Adalatherium hui* (Mammalia, Gondwanatheria) from the Late Cretaceous of Madagascar. Journal of Vertebrate Paleontology, 40(sup1), 133–212.

Hopkins, S. B., & Edward Byrd Davis. (2009). Quantitative morphological proxies for fossoriality in small mammals. Journal of Mammalogy, 90, 1449–60.

Horovitz, I. (2000). The tarsus of Ukhaatherium nessovi (Eutheria, Mammalia) from the Late Cretaceous of Mongolia: an appraisal of the evolution of the ankle in basal therians. Journal of Vertebrate Paleontology, 20(3), 547–560.

Hughes, J. J., Berv, J. S., Chester, S. G., Sargis, E. J., & Field, D. J. (2021). Ecological selectivity and the evolution of mammalian substrate preference across the K–Pg boundary. Ecology and Evolution, 11(21), 14540–14554.

Huttenlocker, A. K., Grossnickle, D. M., Kirkland, J. I., Schultz, J. A., & Luo, Z. X. (2018). Late-surviving stem mammal links the lowermost Cretaceous of North America and Gondwana. Nature, 558(7708), 108–112.

Huxley, T. H. (1880). Arboreal ancestry of the marsupials. In Proceedings of the Zoological Society of London (pp. 655–668).

Jäger, K. R. K., Luo, Z. X., & Martin, T. (2020). Postcranial skeleton of *Henkelotherium guimarotae* (Cladotheria, Mammalia) and locomotor adaptation. Journal of Mammalian Evolution, 27(3), 349–372.

Janis, C. M., & Martín-Serra, A. (2020). Postcranial elements of small mammals as indicators of locomotion and habitat. PeerJ, 8, e9634.

Jenkins Jr, F.A. (1971). The postcranial skeleton of African cynodonts: problems in the early evolution of the mammalian postcranial skeleton. Bulletin of the Peabody Museum of Natural History 36 (1971): 1–216

Jenkins, F.A., Jr. (1974) Tree shrew locomotion and the origins of primate arborealism. In: Primate Locomotion (ed. F.A. Jenkins, Jr.), pp. 85–115. New York: Academic Press.

Jenkins Jr, F. A., & Krause, D. W. (1983). Adaptations for climbing in North American multituberculates (Mammalia). Science, 220(4598), 712–715.

Jenkins, Farish A., & McClearn, D. (1984). “Mechanisms of Hind Foot Reversal in Climbing Mammals.” Journal of Morphology, 182, 197–219.

Jenkins, Jr, F.A., & Parrington, F.R. (1976). The postcranial skeletons of the Triassic mammals *Eozostrodon*, *Megazostrodon*, and *Erythrotherium*. *Philosophical Transactions of the Royal Society of London. B*, Biological Sciences, 273(926), pp.387–431.

Jenkins Jr, F. A., & Schaff, C. R. (1988). The Early Cretaceous mammal *Gobiconodon* (Mammalia, Triconodonta) from the Cloverly Formation in Montana. Journal of Vertebrate Paleontology, 8(1), 1–24.

Ji, Q., Luo, Z. X., Yuan, C. X., Wible, J. R., Zhang, J. P., & Georgi, J. A. (2002). The earliest known eutherian mammal. Nature, 416(6883), 816–822.

Ji, Q., Luo, Z.X., Yuan, C.X., & Tabrum, A.R. (2006). A swimming mammaliaform from the Middle Jurassic and ecomorphological diversification of early mammals. Science, 311(5764), pp.1123–1127.

Jiang, Z. K., Wang, Y. D., Tian, N., Xie, A. W., Zhang, W., Li, L. Q., & Huang, M. (2019). The Jurassic fossil wood diversity from western Liaoning, NE China. Journal of Palaeogeography, 8, 1–11.

Johnson, K.R., & Ellis, B. (2002). A tropical rainforest in Colorado 1.4 million years after the Cretaceous-Tertiary boundary. Science, 296(5577), pp.2379–2383.

Jones, K. E., Bielby, J., Cardillo, M., Fritz, S. A., O’Dell, J., Orme, C. D. L., …, & Purvis, A. (2009). PanTHERIA: a species-level database of life history, ecology, and geography of extant and recently extinct mammals: Ecological Archives E090-184. Ecology, 90(9), 2648–2648.

Jungers, W. L., Falsetti, A. B., & Wall, C. E. (1995). Shape, relative size, and size-adjustments in morphometrics. American Journal of Physical Anthropology, 38(S21), 137–161.

Kemp, T.S. (2005). The Origin and Evolution of Mammals. Oxford University Press.

Kielan-Jaworowska, Z. (1978). Evolution of the therian mammals in the Late Cretaceous of Asia. Part III. Postcranial skeleton in Zalambdalestidae. Palaeontologia Polonica, 38, pp.3–41.

Kielan-Jaworowska, Z., & Gambaryan, P. P. (1994). Postcranial Anatomy and Habits of Asian Multituberculate Mammals. Postcranial Anatomy and Habits of Asian Multituberculate Mammals, 36.

Kielan-Jaworowska, Z., & Qi, T. (1990). Fossorial adaptations of a taeniolabidoid multituberculate mammal from the Eocene of China. Gujizhui dongwu xuebao, 28(2), 83–94.

Kilbourne, B. M. (2017). Selective regimes and functional anatomy in the mustelid forelimb: diversification toward specializations for climbing, digging, and swimming. Ecology and Evolution, 7(21), 8852–8863.

King, B., & Beck, R. M. (2020). Tip dating supports novel resolutions of controversial relationships among early mammals. Proceedings of the Royal Society B, 287(1928), 20200943.

Kirk, E. C., Lemelin, P., Hamrick, M. W., Boyer, D. M., & Bloch, J. I. (2008). Intrinsic hand proportions of euarchontans and other mammals: implications for the locomotor behavior of plesiadapiforms. Journal of Human Evolution, 55(2), 278–299.

Klingenberg, C. P. (2016). Size, shape, and form: concepts of allometry in geometric morphometrics. Development Genes and Evolution, 226(3), 113–137.

Krause, D. W., Hoffmann, S., Hu, Y., Wible, J. R., Rougier, G. W., Kirk, E. C., …, & Rahantarisoa, L. J. (2020). Skeleton of a Cretaceous mammal from Madagascar reflects long-term insularity. Nature, 581(7809), 421–427.

Krause, D. W., & Jenkins, F. A. (1983). The postcranial skeleton of North American multituberculates. Museum of Comparative Zoology, Harvard University.

Lauder, G. V. (1995). On the Inference of Function from Structure. In Functional Morphology in Vertebrate Paleontology, edited by JJ Thomason, 1–18. Cambridge, U.K.: Cambridge University Press.

Lemelin, P. (1999). Morphological correlates of substrate use in didelphid marsupials: implications for primate origins. Journal of Zoology, 247(2), 165–175.

Li, Y., Chang, S. C., Zhang, H., Wang, J., Pei, R., Zheng, D., …, & Hemming, S. R. (2023). A chronostratigraphic and biostratigraphic framework for the Yanliao Biota of northeastern China: Implications for Jurassic terrestrial ecosystems and evolution. Palaeogeography, Palaeoclimatology, Palaeoecology, 630, 111818.

Lungmus, J. K., & Angielczyk, K. D. (2019). Antiquity of forelimb ecomorphological diversity in the mammalian stem lineage (Synapsida). Proceedings of the National Academy of Sciences, 116(14), 6903–6907.

Luo, Z. X. (2007). Transformation and diversification in early mammal evolution. Nature, 450(7172), 1011–1019.

Luo, Z.X., & Wible, J.R. (2005). A Late Jurassic digging mammal and early mammalian diversification. Science, 308(5718), pp.103–107.

Luo, Z. X., Ji, Q., Wible, J. R., & Yuan, C. X. (2003). An Early Cretaceous tribosphenic mammal and metatherian evolution. Science, 302(5652), 1934–1940.

Luo, Zhe-Xi, Chong-Xi Yuan, Qing-Jin Meng, & Qiang Ji. (2011). A Jurassic Eutherian Mammal and Divergence of Marsupials and Placentals. Nature 476: 442–45.

Luo, Z. X., Meng, Q. J., Ji, Q., Liu, D., Zhang, Y. G., & Neander, A. I. (2015). Evolutionary development in basal mammaliaforms as revealed by a docodontan. Science, 347(6223), 760–764.

Luo, Z., Meng, Q., Di, L., Zhang, Y. G., & Yuan, C. X. (2016). Cruro-pedal structure of the paulchoffatiid *Rugosodon eurasiaticus* and evolution of the multituberculate ankle. Palaeontologica Polonica, 67, 149–169.

Luo, Z. X., Meng, Q. J., Grossnickle, D. M., Liu, D., Neander, A. I., Zhang, Y. G., & Ji, Q. (2017). New evidence for mammaliaform ear evolution and feeding adaptation in a Jurassic ecosystem. Nature, 548(7667), 326–329.

MacLeod, N., & Rose, K. D. (1993). Inferring locomotor behavior in Paleogene mammals via eigenshape analysis. American Journal of Science, 293, 300–355.

Malcolm, J.R. (1995). Forest structure and the abundance and diversity of neo-tropical small mammals. In: M.D. Lowman, & N.M. Nadkarni (Eds.). Forest Canopies (pp. 179–197). San Diego, USA: Academic Press.

Mao, F. Y., Zheng, X. T., & Wang, X. L. (2019). Evidence of diphyodonty and heterochrony for dental development in euharamiyidan mammals from Jurassic Yanliao Biota. Vertebrata PalAsiatica, 57(1), 51–76.

Mao, F., Hu, Y., Li, C., Wang, Y., Chase, M. H., Smith, A. K., & Meng, J. (2020). Integrated hearing and chewing modules decoupled in a Cretaceous stem therian mammal. Science, 367(6475), 305–308.

Mao, F., Zhang, C., Liu, C., & Meng, J. (2021). Fossoriality and evolutionary development in two Cretaceous mammaliamorphs. Nature, 592(7855), pp.577–582.

Martin, T. (2005). Postcranial anatomy of *Haldanodon exspectatus* (Mammalia, Docodonta) from the Late Jurassic (Kimmeridgian) of Portugal and its bearing for mammalian evolution. Zoological Journal of the Linnean Society, 145(2), 219–248.

Martin, T., Marugán-Lobón, J., Vullo, R., Martín-Abad, H., Luo, Z. X., & Buscalioni, A. D. (2015). A Cretaceous eutriconodont and integument evolution in early mammals. Nature, 526(7573), 380–384.

Matthew, W. D. (1904). The arboreal ancestry of the Mammalia. The American Naturalist, 38(455/456), 811–818.

Matthew, W. D. (1937). Paleocene faunas of the San Juan Basin, New Mexico. Transactions of the American Philosophical Society, 30, i.

Maynard Smith, J., & Savage, R. J. (1956). Some locomotory adaptations in mammals. Zoological Journal of the Linnean Society, 42(288), 603–622.

McElreath, R. (2020) Statistical Rethinking. 2nd ed. Texts in Statistical Science Series. Boca Raton, Florida, USA: Taylor & Francis Group.

Meng, J. (2014). Mesozoic mammals of China: implications for phylogeny and early evolution of mammals. National Science Review, 1(4), 521–542.

Meng, J., Bi, S., Wang, Y., Zheng, X., & Wang, X. (2014). Dental and mandibular morphologies of *Arboroharamiya* (Haramiyida, Mammalia): a comparison with other haramiyidans and *Megaconus* and implications for mammalian evolution. PloS One, 9(12), e113847.

Meng, J., Hu, Y., Wang, Y., Wang, X., & Li, C. (2006). A Mesozoic gliding mammal from northeastern China. Nature, 444(7121), 889–893.

Meng, Q. J., Grossnickle, D. M., Liu, D., Zhang, Y. G., Neander, A. I., Ji, Q., & Luo, Z. X. (2017). New gliding mammaliaforms from the Jurassic. Nature, 548(7667), 291–296.

Meng, Q.J., Ji, Q., Zhang, Y.G., Liu, D., Grossnickle, D.M., & Luo, Z.X. (2015). An arboreal docodont from the Jurassic and mammaliaform ecological diversification. Science, 347(6223), 764–768.

Merritt, J.F. (2010). The biology of small mammals. Johns Hopkins University Press.

Mosimann, J. E. (1970). Size allometry: size and shape variables with characterizations of the lognormal and generalized gamma distributions. Journal of the American Statistical Association, 65(330), 930–945.

Nations, J. A., Heaney, L. R., Demos, T. C., Achmadi, A. S., Rowe, K. C., & Esselstyn, J. A. (2019). A simple skeletal measurement effectively predicts climbing behaviour in a diverse clade of small mammals. Biological Journal of the Linnean Society, 128(2), 323–336.

Nice, M. M., Nice, C., & Ewers, D. (1956). Comparison of behavior development in snowshoe hares and red squirrels. Journal of Mammalogy, 37(1), 64–74.

Nitikman L.Z., & Mares M.A. (1987). Ecology of small mammals in a gallery forest of central Brazil. Annals of the Carnegie Museum, 56:75–95.

Nowak, R. M. (1991). Walker’s Mammals of the World. 5th ed. Vol. 1. Johns Hopkins Press.

O’Leary, M. A., Bloch, J. I., Flynn, J. J., Gaudin, T. J., Giallombardo, A., Giannini, N. P., …, & Cirranello, A. L. (2013). The placental mammal ancestor and the post–K-Pg radiation of placentals. Science, 339(6120), 662–667.

Panciroli, E., Benson, R.B., Fernandez, V., Humpage, M., Martín-Serra, A., Walsh, S., Luo, Z.X., & Fraser, N.C. (2022). Postcrania of *Borealestes* (Mammaliformes, Docodonta) and the emergence of ecomorphological diversity in early mammals. Palaeontology, 65(1), p.e12577.

Pevsner, S. K., Grossnickle, D. M., & Luo, Z. X. (2022). The functional diversity of marsupial limbs is influenced by both ecology and developmental constraint. Biological Journal of the Linnean Society, 135(3), 569–585.

Polly, P. D. (2007). Limbs in mammalian evolution. In Ed. Hall, B. K. Fins into limbs: evolution, development, and transformation, 15, 245–268.

Polly, P. D., Lawing, A. M., Fabre, A. C., & Goswami, A. (2013). Phylogenetic principal components analysis and geometric morphometrics. Hystrix, 24(1), 33–41.

Prasad, G.V.R., & Godinot, M. (1994). Eutherian tarsal bones from the Late Cretaceous of India. Journal of Paleontology, 68(4), 892–902.

Price, S. A., Friedman, S. T., Corn, K. A., Martinez, C. M., Larouche, O., & Wainwright, P. C. (2019). Building a body shape morphospace of teleostean fishes. Integrative and Comparative Biology, 59(3), 716–730.

Romer, A.S. (1922). The locomotor apparatus of certain primitive and mammal-like reptiles. Bulletin of American Museum of Natural History, 46, pp.517–606.

Romer, A.S., Price, L.W., & Price, L.I. (1940). Review of the Pelycosauria (Vol. 28). Geological Society of America.

Rose, K. D. (2006). The beginning of the age of mammals. JHU Press.

Rougier,, G. W., Qiang, J., & Novacek, M. J 2003). A new symmetrodont mammal with fur impressions from the Mesozoic of China. Acta Geologica Sinica-English Edition, 77(1), 7–14.

Rougier, G. W., Wible, J. R., Beck, R. M., & Apesteguía, S. (2012). The Miocene mammal *Necrolestes* demonstrates the survival of a Mesozoic nontherian lineage into the late Cenozoic of South America. Proceedings of the National Academy of Sciences, 109(49), 20053–20058.

Rowe, T.B., & Greenwald, N.S. (1987). The phylogenetic position and origin of Multituberculata. Journal of Vertebrate Paleontology, 7, pp.24A–25A.

Runestad, J. A., & Ruff, C. B. (1995). Structural adaptations for gliding in mammals with implications for locomotor behavior in paromomyids. American Journal of Physical Anthropology, 98(2), 101–119.

Salton, J. A., & Sargis, E. J. (2008). Evolutionary morphology of the Tenrecoidea (Mammalia) carpal complex. Biological Journal of the Linnean Society, 93, 267–288.

Salton, J. A., & Sargis, E. J. (2009). Evolutionary morphology of the Tenrecoidea (Mammalia) hindlimb skeleton. Journal of Morphology, 270(3), 367–387.

Samuels, J. X., Meachen, J. A., & Sakai, S. A. (2013). Postcranial morphology and the locomotor habits of living and extinct carnivorans. Journal of Morphology, 274(2), 121–146.

Samuels, J. X., & Van Valkenburgh, B. (2008). Skeletal indicators of locomotor adaptations in living and extinct rodents. Journal of Morphology, 269(11), 1387–1411.

Sargis, E. J. (2002a). Functional morphology of the forelimb of tupaiids (Mammalia, Scandentia) and its phylogenetic implications. Journal of Morphology, 253(1), 10–42.

Sargis, E.J. (2002b). Functional morphology of the hindlimb of tupaiids (Mammalia, Scandentia) and its phylogenetic implications. Journal of Morphology, 254(2), 149–185.

Sargis, E. J., Boyer, D. M., Bloch, J. I., & Silcox, M. T. (2007). Evolution of pedal grasping in Primates. Journal of Human Evolution, 53(1), 103–107.

Schmitz, L., & Motani, R. (2011). Nocturnality in dinosaurs inferred from scleral ring and orbit morphology. Science, 332(6030), 705–708.

Shattuck M.R., Williams S.A. (2010). Arboreality has allowed for the evolution of increased longevity in mammals. Proceedings of the National Academy of Science 107:4635–4639.

Shelley SL, Brusatte SL, Williamson TE. (2021) Quantitative assessment of tarsal morphology illuminates locomotor behaviour in Palaeocene mammals following the end-Cretaceous mass extinction. Proceedings of the Royal Society B, 288(20210393).

Silcox, M.T., Bloch, J.I., Boyer, D.M., Chester, S.G., & López-Torres, S. (2017). The evolutionary radiation of plesiadapiforms. *Evolutionary Anthropology: Issues*, News, and Reviews, 26(2), 74–94.

Silcox, M. T., Gunnell, G. F. (2008). Plesiadapiformes. In C. M. Janis, G. F. Gunnell, & M. D. Uhen (Eds.) Evolution of Tertiary Mammals of North America Vol. 2: Small Mammals, Xenarthrans, and Marine Mammals (pp. 207–238). Cambridge, UK: Cambridge University Press.

Simpson, G. G. (1926). Mesozoic Mammalia. IV. The multituberculates as living animals. American Journal of Science, 11(63), 228–250.

Simpson, G. G., & Elftman, H. O. (1928). Hind limb musculature and habits of a Paleocene multituberculate. American Museum novitates; no. 333.

Spindler, F., Werneburg, R., Schneider, J.W., Luthardt, L., Annacker, V., & Rößler, R., 2018. First arboreal ‘pelycosaurs’ (Synapsida: Varanopidae) from the early Permian Chemnitz Fossil Lagerstätte, SE Germany, with a review of varanopid phylogeny. PalZ, 92, 315–364.

Sues, H.D., & Jenkins, F.A. (2006). The postcranial skeleton of *Kayentatherium wellesi* from the Lower Jurassic Kayenta Formation of Arizona and the phylogenetic significance of postcranial features. Amniote paleobiology: perspectives on the evolution of mammals, birds, and reptiles, 114.

Sumida, S.S., & Modesto, S. (2001). A phylogenetic perspective on locomotory strategies in early amniotes. American Zoologist, 41(3), 586–597.

Sussman, R. W. (1991). Primate origins and the evolution of angiosperms. American Journal of Primatology, 23(4), 209–223.

Szalay, F.S., 1981. Functional analysis and the practice of the phylogenetic method as reflected by some mammalian studies. American Zoologist, 21(1), pp.37–45.

Szalay, F. S. (1984). Arboreality: is it homologous in metatherian and eutherian mammals?. In Evolutionary Biology: Volume 18 (pp. 215–258). Boston, MA: Springer US.

Szalay, F. S. (1994). Evolutionary history of the marsupials and an analysis of osteological characters. Cambridge University Press.

Szalay, F.S., & Decker, R.L. (1974). Origins, evolution, and function of the tarsus in Late Cretaceous Eutheria and Paleocene primates. Primate locomotion, 1, 223–259.

Szalay, F. S., & Sargis, E. J. (2001). Model-based analysis of postcranial osteology of marsupials from the Palaeocene of Itaboraí (Brazil) and the phylogenetics and biogeography of Metatheria. Geodiversitas, 23(2), 139–302.

Szalay, F. S., & Sargis, E. J. (2006). Cretaceous therian tarsals and the metatherian-eutherian dichotomy. Journal of Mammalian Evolution, 13, 171–210.

Szalay, F. S., & Trofimov, B. A. (1996). The Mongolian Late Cretaceous Asiatherium, and the early phylogeny and paleobiogeography of Metatheria. Journal of Vertebrate Paleontology, 16(3), 474–509.

Thorington Jr, R. W., Miller, A. M., & Anderson, C. G. (1998). Arboreality in tree squirrels (Sciuridae). In Ecology and evolutionary biology of tree squirrels (Vol. 6, pp. 119–130). Virginia Museum of Natural History.

Trofimov, B. A., & Szalay, F. S. (1994). New Cretaceous marsupial from Mongolia and the early radiation of Metatheria. Proceedings of the National Academy of Sciences, 91(26), 12569–12573.

Upham, N. S., Esselstyn, J. A., & Jetz, W. (2019). Inferring the mammal tree: species-level sets of phylogenies for questions in ecology, evolution, and conservation. PLoS biology, 17(12), e3000494.

Urbani, B., & Youlatos, D. (2013). Positional behavior and substrate use of *Micromys minutus* (Rodentia: Muridae): insights for understanding primate origins. Journal of Human Evolution, 64(2), 130–136.

Vehtari, A., Gelman, A., & Gabry, J. (2017). Practical Bayesian model evaluation using leave-one-out cross-validation and WAIC. Statistics and computing, 27, 1413–1432.

Wagner, B., Kreft, H., Nitschke, C. R., & Schrader, J. (2023). Remotely sensed tree height and density explain global gliding vertebrate richness. Ecography, e06435.

Wang, H. B., Hoffmann, S., Wang, D. C., & Wang, Y. Q. (2022). A new mammal from the Lower Cretaceous Jehol Biota and implications for eutherian evolution. Philosophical Transactions of the Royal Society B, 377(1847), 20210042.

Wainwright, Peter C., Michael E. Alfaro, Daniel I. Bolnick, & C. Darrin Hulsey. (2005). Many-to-One Mapping of Form to Function: A General Principle in Organismal Design? Integrative and Comparative Biology, 45(2), 256–262.

Wang, H., Meng, J., & Wang, Y. (2019). Cretaceous fossil reveals a new pattern in mammalian middle ear evolution. Nature, 576(7785), 102–105.

Weaver, L. N., & Grossnickle, D. M. (2020). Functional diversity of small-mammal postcrania is linked to both substrate preference and body size. Current zoology, 66(5), 539–553.

Weaver, L. N., Varricchio, D. J., Sargis, E. J., Chen, M., Freimuth, W. J., & Wilson Mantilla, G. P. (2021). Early mammalian social behaviour revealed by multituberculates from a dinosaur nesting site. Nature Ecology & Evolution, 5(1), 32–37.

Weisbecker, V., & D. I. Warton. (2006). Evidence at Hand: Diversity, Functional Implications, and Locomotor Prediction in Intrinsic Hand Proportions of Diprotodontian Marsupials. Journal of Morphology, 267, 1469–85.

Weisbecker, V., & Schmid, S. (2007). Autopodial skeletal diversity in hystricognath rodents: functional and phylogenetic aspects. Mammalian Biology, 72(1), 27–44.

Williamson, T. E., Brusatte, S. L., & Wilson, G. P. (2014). The origin and early evolution of metatherian mammals: the Cretaceous record. ZooKeys*, (*465), 1.

Wilman, H., Belmaker, J., Simpson, J., de la Rosa, C., Rivadeneira, M. M., & Jetz, W. (2014). EltonTraits 1.0: Species-level foraging attributes of the world’s birds and mammals: Ecology, 95(7), 2027–2027.

Wilson Mantilla, G. P., Chester, S. G., Clemens,W. A., Moore, J. R., Sprain, C. J., Hovatter, B. T., …, & Renne, P. R. (2021). Earliest Palaeocene purgatoriids and the initial radiation of stem primates. Royal Society Open Science, 8(2), 210050.

Wing, S.L., & Sues, H.D. (1992). Mesozoic and early Cenozoic terrestrial ecosystems. Terrestrial ecosystems through time: evolutionary paleoecology of terrestrial plants and animals.

Wing, S.L., & Tiffney, B.H. (1987). The reciprocal interaction of angiosperm evolution and tetrapod herbivory. Review of Palaeobotany and Palynology, 50(1-2), 179–210.

Woodman, N., & Stabile, F. A. (2015). Functional skeletal morphology and its implications for locomotory behavior among three genera of myosoricine shrews (Mammalia: Eulipotyphla: Soricidae). Journal of Morphology, 276(5), 550–563.

Woodman, N. (2023). Skeletal indicators of locomotor adaptations in shrews. Therya, 14(1), 15–37.

Wu, J., Yonezawa, T., & Kishino, H. (2017). Rates of molecular evolution suggest natural history of life history traits and a post-K-Pg nocturnal bottleneck of placentals. Current Biology, 27(19), 3025–3033.

Xu, X., Zhou, Z., Sullivan, C., Wang, Y., & Ren, D. (2016). An updated review of the Middle-Late Jurassic Yanliao Biota: Chronology, taphonomy, paleontology and paleoecology. Acta Geologica Sinica-English Edition, 90(6), 2229–2243.

Youlatos, D., Karantanis, N. E., Byron, C. D., & Panyutina, A. (2015). Pedal grasping in an arboreal rodent relates to above-branch behavior on slender substrates. Journal of Zoology, 296(4), 239–248.

Yuan, C.X., Ji, Q., Meng, Q.J., Tabrum, A.R., & Luo, Z.X. (2013). Earliest evolution of multituberculate mammals revealed by a new Jurassic fossil. Science, 341(6147), 779–783.

Zelditch, M., Swiderski, D., & Sheets, H. D. (2012). Geometric morphometrics for biologists: a primer. Academic Press.

Zhou, C. F., Wu, S., Martin, T., & Luo, Z. X. (2013). A Jurassic mammaliaform and the earliest mammalian evolutionary adaptations. Nature, 500(7461), 163–167.

