## Supplementary figures and images for "Locomotor Trends Among Early Mammals Illuminated by Predictors of Arboreality from the Mammalian Appendicular Skeleton"

### Figures_S1-S3

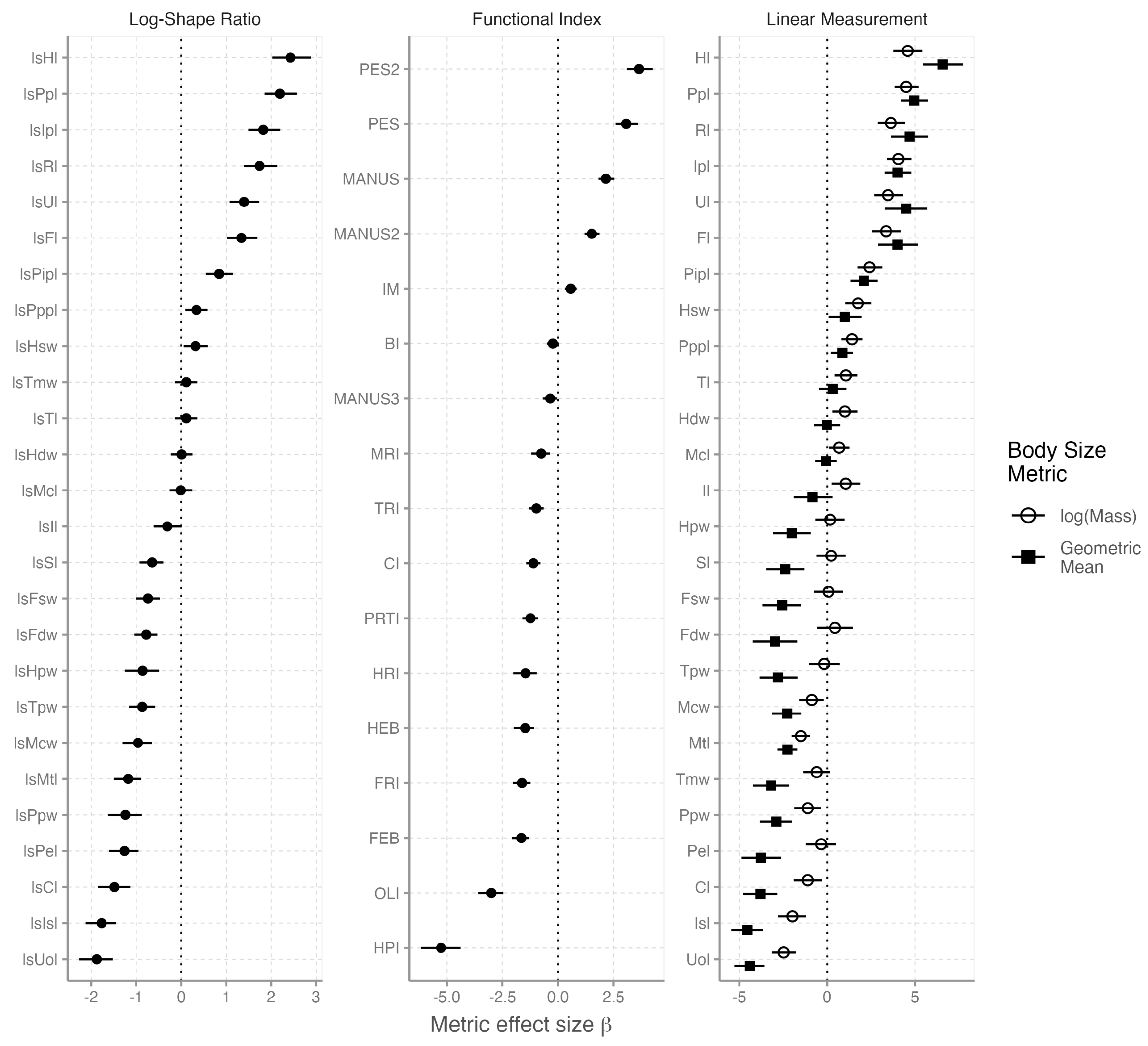

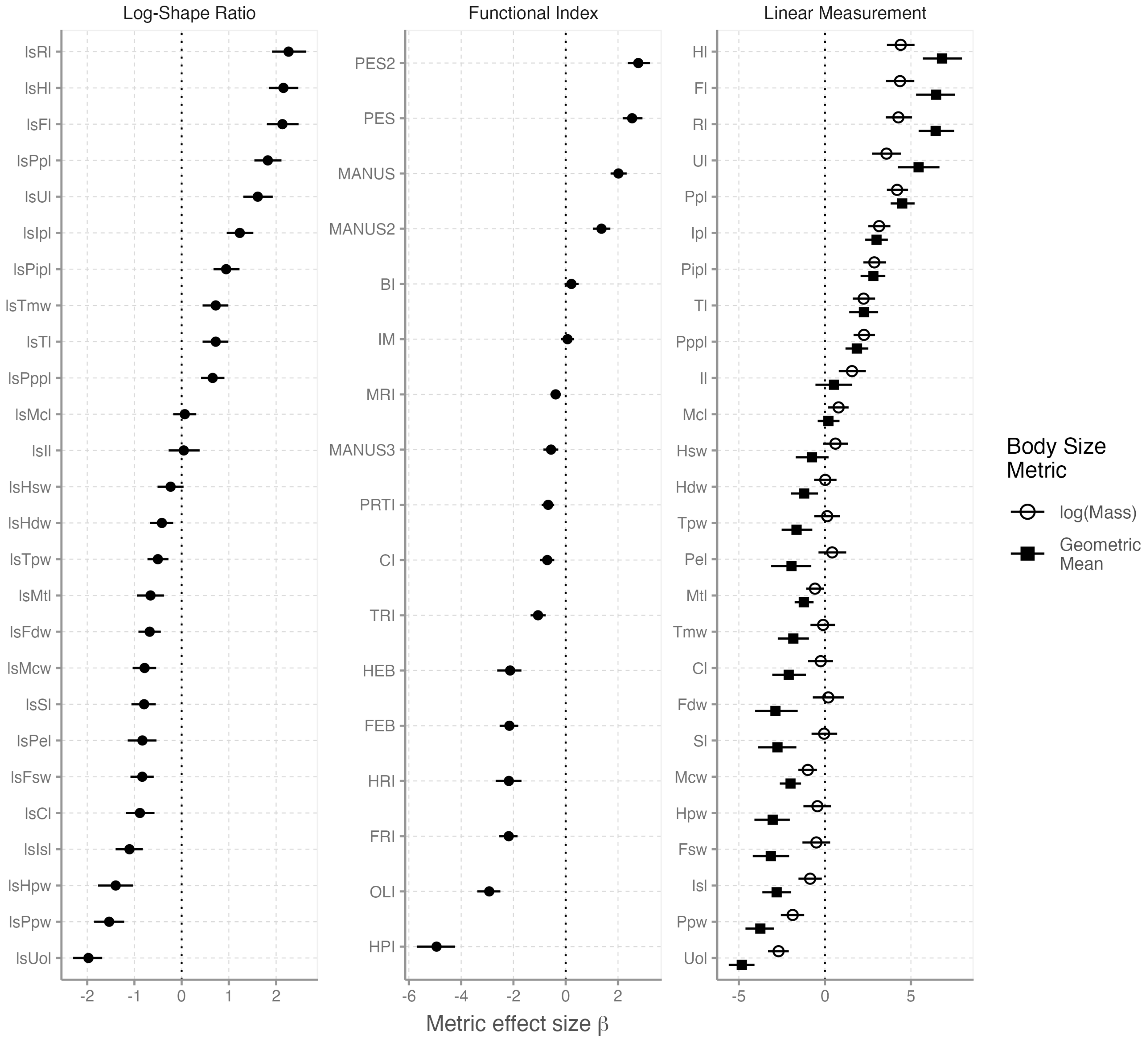

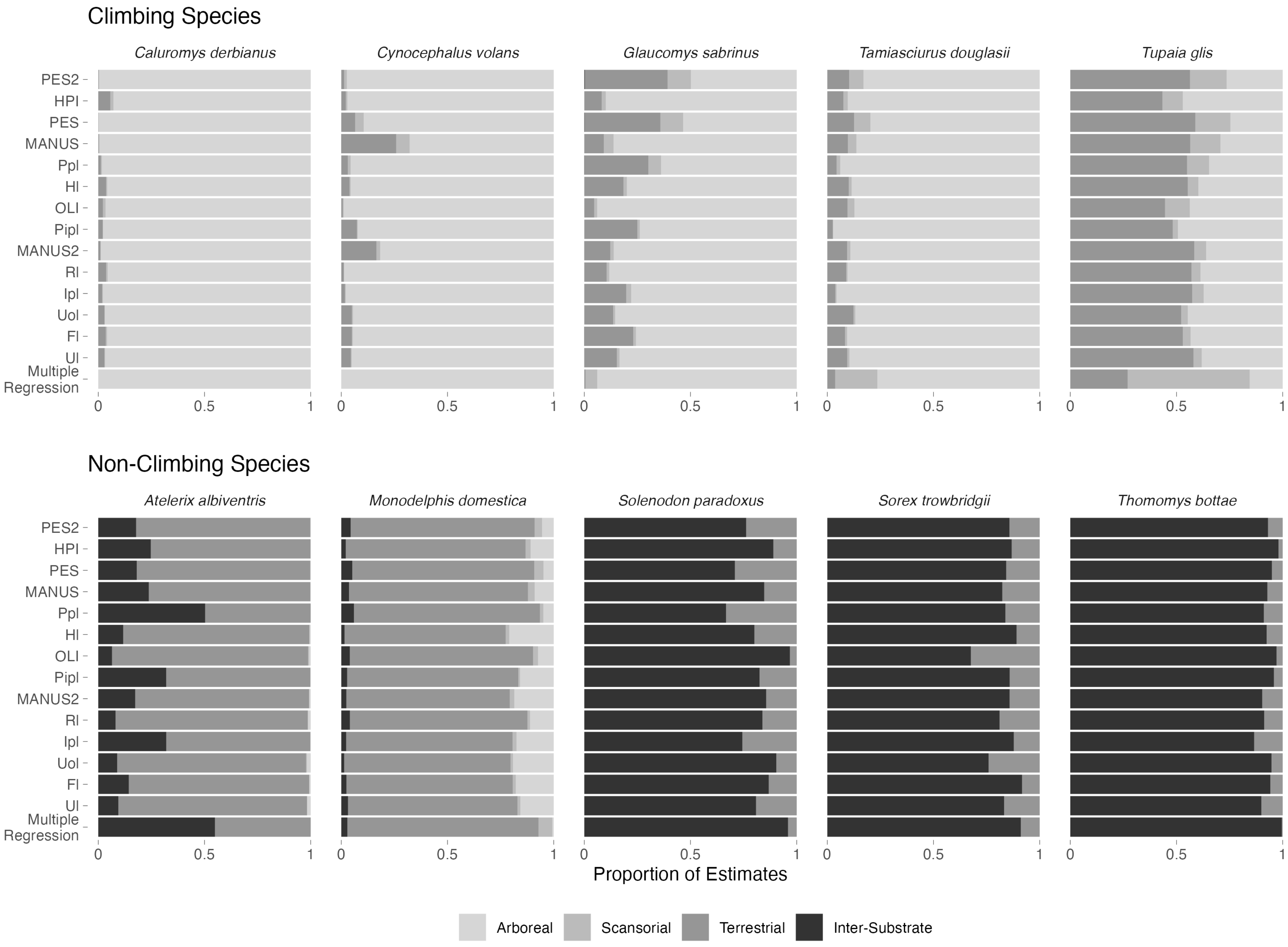
